# Host and antibiotic jointly select for greater virulence in *Staphylococcus aureus*

**DOI:** 10.1101/2024.08.31.610628

**Authors:** Michelle Su, Kim L. Hoang, McKenna Penley, Michelle H. Davis, Jennifer D. Gresham, Levi T. Morran, Timothy D. Read

## Abstract

Widespread antibiotic usage has resulted in the rapid evolution of drug-resistant bacterial pathogens and poses significant threats to public health. Resolving how pathogens respond to antibiotics under different contexts is critical for understanding disease emergence and evolution going forward. The impact of antibiotics has been demonstrated most directly through *in vitro* pathogen passaging experiments. Independent from antibiotic selection, interactions with hosts have also altered the evolutionary trajectories and fitness landscapes of pathogens, shaping infectious disease outcomes. However, it is unclear how interactions between hosts and antibiotics impact the evolution of pathogen virulence. Here, we evolved and re-sequenced *Staphylococcus aureus,* a major bacterial pathogen, varying exposure to host and antibiotics to tease apart the contributions of these selective pressures on pathogen adaptation. After 12 passages, *S. aureus* evolving in *Caenorhabditis elegans* nematodes exposed to a sub-minimum inhibitory concentration of antibiotic (oxacillin) became highly virulent, regardless of whether the ancestral pathogen was methicillin-resistant (MRSA) or methicillin-sensitive (MSSA). Host and antibiotic exposure selected for reduced drug susceptibility in MSSA lineages while increasing MRSA total growth outside hosts. We identified mutations in genes involved in complex regulatory networks linking virulence and metabolism, including *codY*, *agr*, and *gdpP*, suggesting that rapid adaptation to infect hosts may have pleiotropic effects. In particular, MSSA populations under selection from host and antibiotic accumulated mutations in the global regulator gene *codY*, which controls biofilm formation in *S. aureus.* These populations had indeed evolved more robust biofilms—a trait linked to both virulence and antibiotic resistance—suggesting evolution of one trait can confer multiple adaptive benefits. Mutations that arose in these genes were also enriched in clinical isolates associated with systemic infections in humans. Despite evolving in similar environments, MRSA and MSSA populations—differing only in the presence of an intact accessory gene (*mecA)*—proceeded on divergent evolutionary paths, with MSSA populations exhibiting more similarities across replicate populations. Our results underscore the importance of considering the host context as a critical driver of pathogen traits like virulence and antibiotic resistance.

## INTRODUCTION

Antibiotic resistance is a global crisis (Larsson and Flach, 2022). Pathogens that evolve resistance and are able to survive antibiotic therapy can go on to cause disease and transmit to other hosts; hence antibiotic resistance and virulence are intertwined (Geisinger and Isberg, 2017). Selective pressures like antibiotic usage and host defenses have independently been shown to alter the evolution of pathogen traits (Herren and Baym, 2022; Kim L Hoang et al., 2024; Kubinak et al., 2012; Larsson and Flach, 2022; Toprak et al., 2012). However, pathogens face multiple selective pressures in their environment, and selection by these forces can interact synergistically to alter evolutionary rates and trajectories relative to individual pressures alone (Merlo et al., 2020; Sharma et al., 2020). It is unclear how virulence evolves in the face of more than one selective pressure and whether this trait is constrained or facilitated by antibiotic resistance. These dynamics are even less understood in the early stages of disease emergence, where pathogens face a different suite of selection dynamics than pathogens having reached equilibrium (Day et al., 2022; Visher et al., 2021). Identifying conditions under which antibiotic resistance and virulence evolve in novel host populations will shed light on how diseases emerge and potential mitigation measures.

The strength of selection impacts pathogen evolution (Cisneros-Mayoral et al., 2022; Mahrt et al., 2021; Oz et al., 2014; Wistrand-Yuen et al., 2018). While antibiotic therapy administers concentrations that should kill all pathogen cells, low concentrations of antibiotics (sub-minimum inhibitory concentration, or sub-MIC) are pervasive in a variety of settings, including in natural environments (e.g., water bodies and soil) due to pollution and biological waste (Larsson and Flach, 2022). Sub-MICs allow susceptible populations to continue dividing, affording opportunities for mutations conferring greater resistance to emerge (Andersson and Hughes, 2014). Ultimately, selection by sub-MICs can lead to highly resistant pathogens (Gullberg et al., 2011; Wistrand-Yuen et al., 2018), which tend to incur less fitness costs compared to those under selection by high antibiotic concentrations (Westhoff et al., 2017). Importantly, mutations not related to antibiotic resistance can arise, such as those involved in adaptation to the growth environment (Pereira et al., 2023). Low concentrations of antibiotics can also alter expression of virulence factors *in vitro* across a broad range of pathogens (Braga et al., 2000; El-Houssaini et al., 2019; Haddadin et al., 2010; Khan et al., 2020), suggesting that they can affect virulence during infection. Sub-MIC antibiotics therefore have the potential to alter the early stages of disease emergence in host populations, particularly for those pathogens that can spend parts of their life history outside the host. Taken together, exposure to low antibiotic concentrations likely alters how pathogens interact with their hosts, shaping evolution of both virulence and antibiotic resistance. However, there is a knowledge gap in the interaction between antibiotics and pathogen evolution *in vivo* (Windels et al., 2020).

The host can exert strong selection on pathogens, especially during the period when a new organism is infected (e.g., a zoonotic transition). Infection of new host individuals tends to bottleneck pathogen populations (Bacigalupe et al., 2019; Klemm et al., 2016). These bottlenecks can confer a fitness advantage to resistant bacteria in the face of low antibiotic concentrations (McVicker et al., 2014). In addition to defenses like the immune system, the host environment differs drastically in nutrient availability compared to rich laboratory media (Windels et al., 2020). For example, *Staphylococcus aureus* passaged through macrophage cell lines had increased pathogen survival as well as resistance to antibiotics; these traits were lost when the pathogen was exposed to nutrient-rich media (Alves et al., 2024). Similarly, the human defensive peptide b-defensin 3 can maintain reduced susceptibility of *S. aureus* to the antibiotic vancomycin (Fait et al., 2023). Adapting to different niches within a host can also bring about drastic genomic changes. *Salmonella enterica* transitioning from an intestinal to a systemic lifestyle exhibited genome degradation in genes no longer necessary for inhabiting the gastrointestinal tract (Klemm et al., 2016). While sub-MIC antibiotic exposure and host factors have been shown to independently shape pathogen evolution, it is unclear how the collective actions of both selective pressures affect the evolution of virulence and antibiotic resistance.

*Staphylococcus aureus* is an opportunistic pathogen that commonly colonizes nares and skin in humans but can invade internal organs and blood, causing systemic infections and bacteremia (Tsouklidis et al., 2020). In 2017, more than 119,000 bacteremia infections caused by *S. aureus* occurred in the United States with a mortality rate of 18% (Kourtis et al., 2019). *S. aureus* colonizing host surfaces exhibit distinct genomic signatures against those isolated during systemic infection, indicating that the pathogen employs different strategies to adapt to different host sites (Giulieri et al., 2022). For example, hemolytic strains tend to be more abundant than non-hemolytic ones in murine systemic infection models, whereas the opposite occurs for wound models (Schwan et al., 2003). Once inside the host, *S. aureus* resists host immune function by hindering or lysing immune cells (Kwiecinski and Horswill, 2020; Thomer et al., 2016). Sub-MIC concentrations of beta-lactam antibiotics—which disrupts bacterial cell wall synthesis—modify expression of virulence factors in *S. aureus*, increasing the expression of alpha-hemolysin (Hodille et al., 2017). Treatment with beta-lactam antibiotics systemically exposes *S. aureus* to low antibiotic levels at non-target host tissues (Nix et al., 1991). As an opportunistic pathogen, *S. aureus* can also survive for extended periods of time outside of hosts and have been isolated from environments such as veterinary clinics and schools, as well as natural and constructed settings such as wastewater and beaches (Loeffler et al., 2005; Roberts et al., 2013; Steadmon et al., 2023; Thapaliya et al., 2017). Combined with the increasing prevalence and concentrations of antibiotics in the environment, these factors likely increase opportunities for exposure of *S. aureus* to sub-MIC antibiotics, subsequently affecting its interaction with the host during infection.

Experimental evolution has been used to elucidate how different selective pressures impact pathogen evolution. *In vitro* studies have yielded insights into the phenotypes and genetic loci generated by longer-term antibiotic selection through passaging experiments lasting hundreds of generations (Fait et al., 2023; Long et al., 2023; Westhoff et al., 2017; Wistrand-Yuen et al., 2018). Conversely, *in vivo* studies have focused on transmitting pathogens through individual hosts for a single to a handful of passages (Erler et al., 2024; Higazy et al., 2024; McVicker et al., 2014). However, few systems have been suitable to examine how antibiotics affect pathogen adaptation within a host context. In this study, we take advantage of the suitability of a multicellular host, *Caenorhabditis elegans* nematodes, to study designs of high replication across an appreciable temporal scale. The *S. aureus* used here was originally isolated from a skin and soft tissue infection of a human inmate (Diep et al., 2006) and thus is a novel pathogen to *C. elegans (Ekroth et al., 2021)*. Nonetheless, *S. aureus* can kill nematodes by colonizing the host intestine and lysing cells, inducing expression of defense genes in nematodes with roles conserved in humans (Irazoqui et al., 2010; Sifri et al., 2003). Virulence screens in *C. elegans* using *S. aureus* transposon libraries also showed overlapping results to other infection models (Bae et al., 2004; Begun et al., 2005), with the degree of *S. aureus* virulence in *C. elegans* reflecting disease severity of human infections (Wu et al., 2013, 2010). Experimentally evolving *S. aureus* in *C. elegans* thus allows us to track the early stages of virulence and antibiotic resistance evolution in novel host populations with the potential to identify conserved genomic regions underlying evolved traits.

Here, we directly test the impact of host and sub-MIC antibiotic exposure on pathogen evolution. Selection exerted by two forces may impede the pathogen’s response to one or both forces (Merlo et al., 2020). Adaptation may require resources to be expended toward either virulence or antibiotic resistance, leading to a trade-off between these traits (Ferenci, 2016). Alternatively, weaker selection from sub-MIC antibiotics may interact synergistically with hosts and facilitate the evolution or maintenance of high virulence and antibiotic resistance. Sub-MIC antibiotics can favor no-cost mutations (Westhoff et al., 2017), wherein pathogens can rapidly adapt without affecting other traits (Visher et al., 2021). Because emerging pathogens tend to be far from the optimum of the fitness landscape, we expect pleiotropic or large-effect mutations to play a role (Bomblies and Peichel, 2022). We took advantage of the tractability of the system to determine how virulence, fitness, and antibiotic resistance are connected and how multiple selective pressures shape the evolutionary trajectory of two *S. aureus* isolates differing only in their antibiotic susceptibility.

Carriage of Staphylococcal cassette chromosome *mec* (SCC*mec*), which encodes *mecA*, an accessory gene that provides resistance against beta-lactam antibiotics, is a major mechanism for existing antibiotic resistance in *S. aureus* (Shore and Coleman, 2013). We passaged two *S. aureus* isogenic strains, one with existing resistance (*mecA+*, MRSA) and one with a transposon insertion in *mecA* (*mecA-*, MSSA) under selection by *C. elegans* and a sub-MIC level of oxacillin, a beta-lactam antibiotic that has replaced methicillin as a therapy for Staphylococcal infections. We selected for virulence in both strains to determine whether antibiotic resistance can hinder the evolution of greater virulence. We then quantified the ability of evolved pathogens to kill hosts as the metric for virulence. We also assessed whether populations maintained their ability to hemolyze red blood cells—an indicator of virulence expression in both humans and nematodes (Sifri et al., 2003). Because growth outside the host is important for transmission of opportunistic pathogens, we quantified the *in vitro* growth and MIC of evolved pathogens. Finally, we identified mutations potentially underlying evolved traits and compared them against a database of over 80,000 *S. aureus* genomes to ascertain whether these mutations may be associated with adaptation to different human host sites.

## RESULTS

### Host and sub-MIC antibiotic exposure selected for greater virulence in both MRSA and MSSA

We experimentally evolved two *S. aureus* isogenic variants of USA300 JE2 (Diep et al., 2006), MRSA and MSSA, with or without a host and sub-MIC antibiotic exposure. The ancestral MRSA and MSSA isolates did not differ in terms of total growth without oxacillin, but exhibited a slight decline in sub-MIC oxacillin (Figures 1A and 1B). For each ancestor, we passaged six independently evolving populations under each condition 12 times, for a total of 48 evolved *S. aureus* populations (Figure 1C).

**Figure 1.**
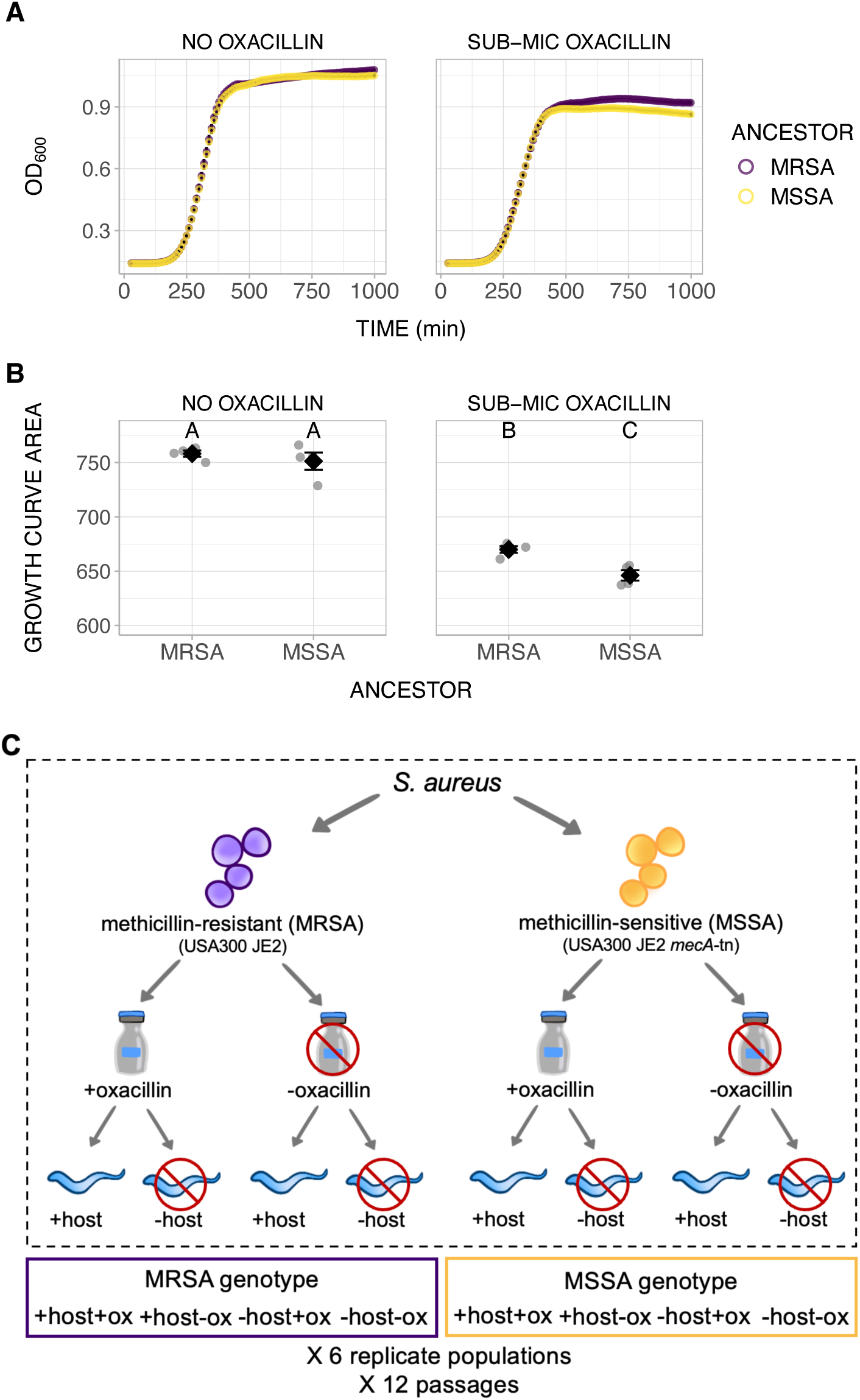
Experimental evolution design. A) Growth curves and B) total growth of ancestral MRSA and MSSA populations *in vitro* with and without sub-MIC oxacillin. Different letters indicate significant differences (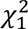 = 6.39, P = 0.01). C) MRSA and MSSA were passaged 12 times with or without hosts, in the presence or absence of a sub-MIC of the antibiotic oxacillin. Each treatment consisted of six independently evolving replicate populations. Experimental evolution treatment abbreviations are indicated in the purple and yellow boxes. Error bars indicate standard errors.

To assess the changes in virulence of evolved *S. aureus*, we measured the mortality of *C. elegans* infected with evolved pathogens. While the MRSA and MSSA ancestors caused similar levels of host mortality (dotted and dashed lines in Figure 2A), there was a significant treatment effect for evolved pathogens (Figure 2A). For the MRSA genotype, host and oxacillin exposure selected for the greatest virulence. Conversely, pathogens evolved in the absence of either pressure exhibited attenuated virulence. These populations also had significantly greater variance compared to those under selection from host and oxacillin, and those under solely host selection. For MSSA, host and oxacillin exposure similarly favored greater virulence over the other three conditions. There was no difference between variances.

**Figure 2.**
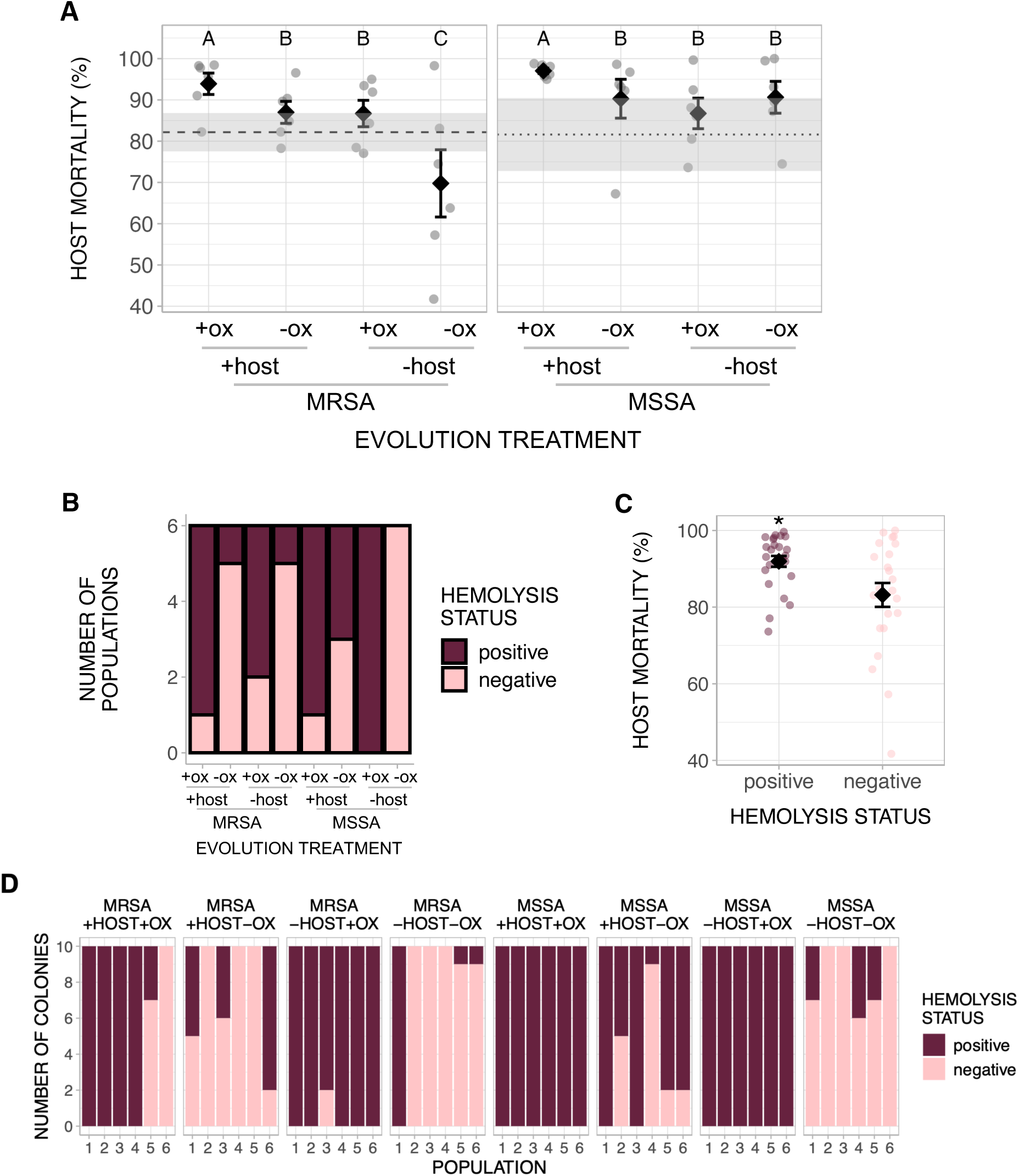
Evolution of virulence is facilitated by exposure to both host and sub-MIC antibiotic. A) Virulence in terms of *C. elegans* mortality. Dashed and dotted lines indicate respective ancestral virulence. Shaded areas indicate standard errors of technical replicates of ancestral virulence. Different letters indicate significant differences (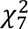 = 460.43, P < 0.001. MRSA Levene’s test for homogeneity of variance: F_3,20_ = 4.06, P = 0.021. MSSA: F_3,20_ = 0.99, P = 0.417). B) Virulence in terms of the ability to hemolyze sheep’s blood, assayed at the population level (oxacillin: 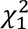 = 5.82, P = 0.016). The y-axis indicates the number of evolved populations for each category. C). Host mortality from A) grouped by hemolysis status in B) (Kruskal-Wallis 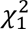 = 3.97, P = 0.046). D) The proportion of colonies sampled from each evolved population that are able to hemolyze sheep’s blood (oxacillin: 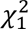 = 23.30, P < 0.001. Levene’s test for homogeneity of variance: F_1,22_ = 26.06, P < 0.001). Error bars indicate standard errors. *P < 0.05

We also characterized the hemolytic activity of evolved populations, which correlates with the ability to secrete extracellular toxins and virulence (Cheung et al., 2012). At least one population from most treatments had lost the ancestral ability to hemolyze sheep’s blood (Figure 2B). Sub-MIC oxacillin maintained hemolytic activity, suggesting that constant exposure to low concentrations of oxacillin favored retention of extracellular toxicity. While hemolysis was not necessary for increased host-killing, this ability was more often found in populations causing greater nematode mortality (Figure 2C). As virulence was still heightened in pathogens unable to destroy red blood cells (e.g., MSSA pathogens evolving without host or oxacillin caused greater than 90% mean mortality despite none being hemolysis-positive), other virulence factors may be compensating for the absence of hemolysis.

Natural populations of *S. aureus* exhibit variation in hemolytic ability (King et al., 2016), even within an individual host (McAdam et al., 2011). We thus hypothesized that a significant selective pressure like oxacillin would favor little variation of this trait within a population. We determined the hemolysis status of isolates from each evolved population (10 isolates x 48 populations)—most populations were in consensus (Figure S1). Furthermore, oxacillin still has a significant effect, where hemolysis was maintained when under sub-MIC exposure selection (Figure 2D). Oxacillin favored less hemolytic diversity in MSSA populations, further demonstrating the interaction between antibiotics and virulence, and that antibiotics can shape variation in virulence traits.

### Host and sub-MIC antibiotic synergistically promoted growth of MRSA outside hosts and reduced drug susceptibility in MSSA

We measured the *in vitro* growth of evolved populations to evaluate how pathogen fitness outside the host had been impacted. We used rich media to replicate the conditions under which *S. aureus* evolved during the experiment. Importantly, rich media reduced the risk of introducing additional selective pressures than those being tested. In media without oxacillin, there were no significant differences in total growth between treatments (Figure 3A). However, in sub-MIC oxacillin, MRSA populations under selection from both pressures exhibited the greatest growth (Figure 3B), with some achieving more growth than those in the absence of oxacillin. By contrast, exposure to sub-MIC oxacillin alone yielded the lowest growth, suggesting a fitness cost. Similarly, MSSA populations exposed to host and sub-MIC oxacillin exhibited a moderate increase in growth compared to other combinations of selective pressures.

**Figure 3.**
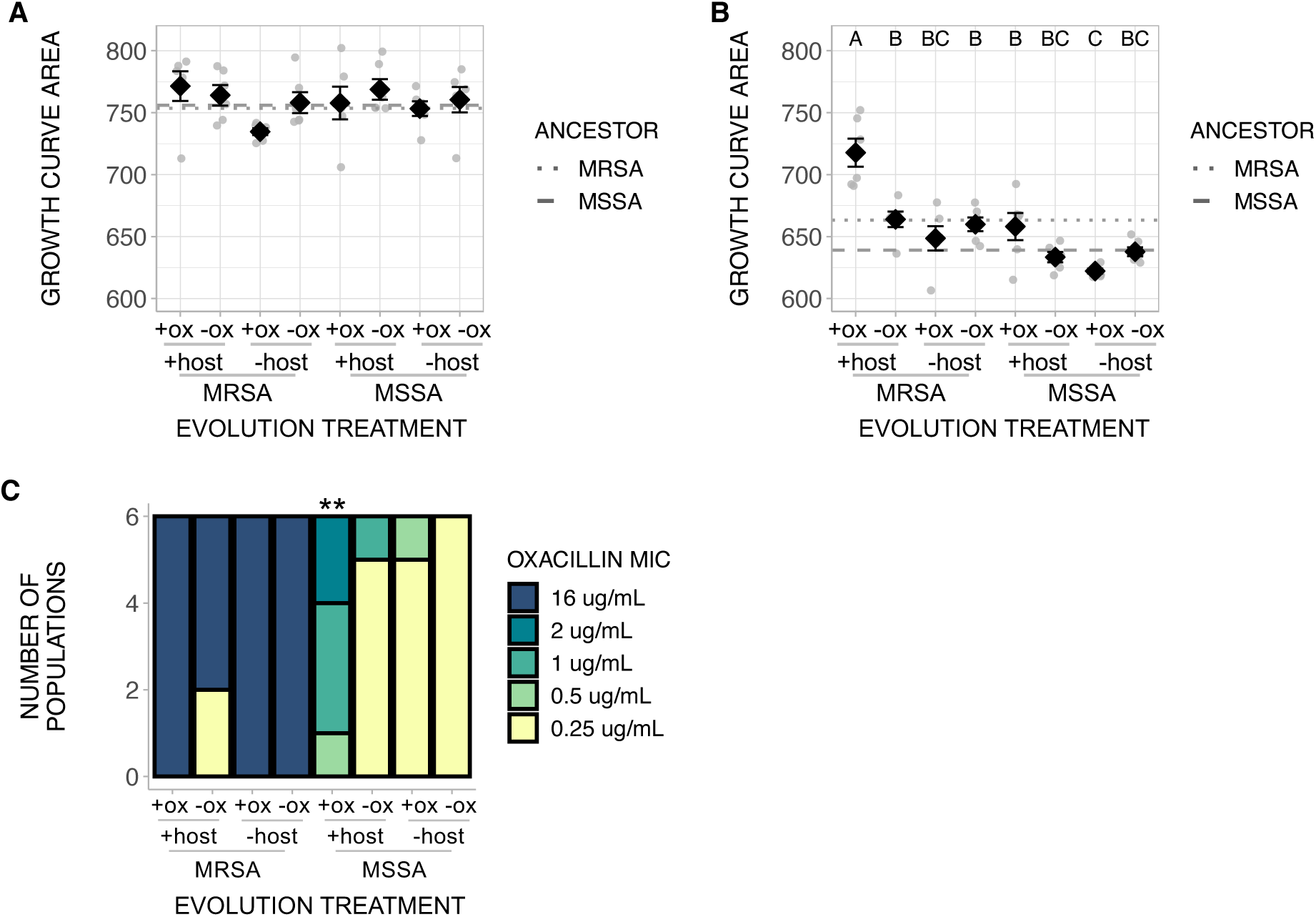
Host and sub-MIC antibiotic selection facilitated pathogen growth in antibiotics. Pathogen *in vitro* growth A) without oxacillin (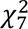 = 9.24, P = 0.24) and B) in sub-MIC oxacillin (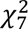 = 106.16, P < 0.001). Different letters indicate significant differences. C). Oxacillin MIC of evolved populations (Fisher’s exact test P < 0.001). The y-axis indicates the number of evolved populations for each category. Error bars indicate standard errors. **P < 0.01

We then determined the MICs of evolved populations to ascertain how host and sub-MIC exposure affected the evolution and maintenance of antibiotic resistance. For the MRSA genotype, all but two populations retained the ancestral level of oxacillin resistance (16ug/mL oxacillin; Figure 3C). During evolution in hosts without antibiotic selection, two populations lost their resistance and exhibited similar susceptibility as the MSSA ancestor (0.25ug/mL). For the MSSA genotype, host and oxacillin exposure selected for decreased antibiotic sensitivity, up to 8-fold the ancestral MIC. These results suggested sub-MIC exposure combined with host factors potentiated the increase in antibiotic resistance in MSSA. One population under selection from solely the host also had an increased MIC, supporting previous evidence showing non-antibiotic selective pressures, such as a host in our study, can select for reduced antibiotic susceptibility (Koch et al., 2014). There was no variation in antibiotic resistance within MRSA and little variation within MSSA across the most genetically diverse populations (Figure S2).

### Parallel evolution of regulators of virulence and antibiotic resistance across evolved populations

We conducted whole-genome sequencing of populations to identify mutations arisen from host and antibiotic selection. Below we focus on mutations that had swept to fixation (Figure 4A, Table S1); Figures S3 and S4 summarize mutations that were below 100% frequency in each population. Populations evolved from the MSSA ancestor had significantly fewer mutations (excluding those intergenic or synonymous) than MRSA populations (Figure S5; 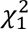 = 5.28, P = 0.022), potentially due to the slightly reduced growth of MSSA in the presence of oxacillin (Figure 1B). Sub-MIC oxacillin selection also resulted in more mutations than in its absence (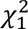 = 5.92, P = 0.015), although this is likely driven by MRSA populations. While this result is consistent with the role of antibiotics in increasing mutation rates in bacteria (Revitt-Mills and Robinson, 2020), there were only two mutations in DNA and mismatch repair genes (*mutL* and *recA*), suggesting repair genes were not the sole mechanism involved. Of the 32 indels detected, 19 were in host-associated populations. Across all populations, many mutations occurred in the same gene (e.g., *agr, gdpP*, *graSR, pbpA,* and *saeRS*) but not the same amino acid (Table S2), indicating parallel evolution at the gene level.

**Figure 4.**
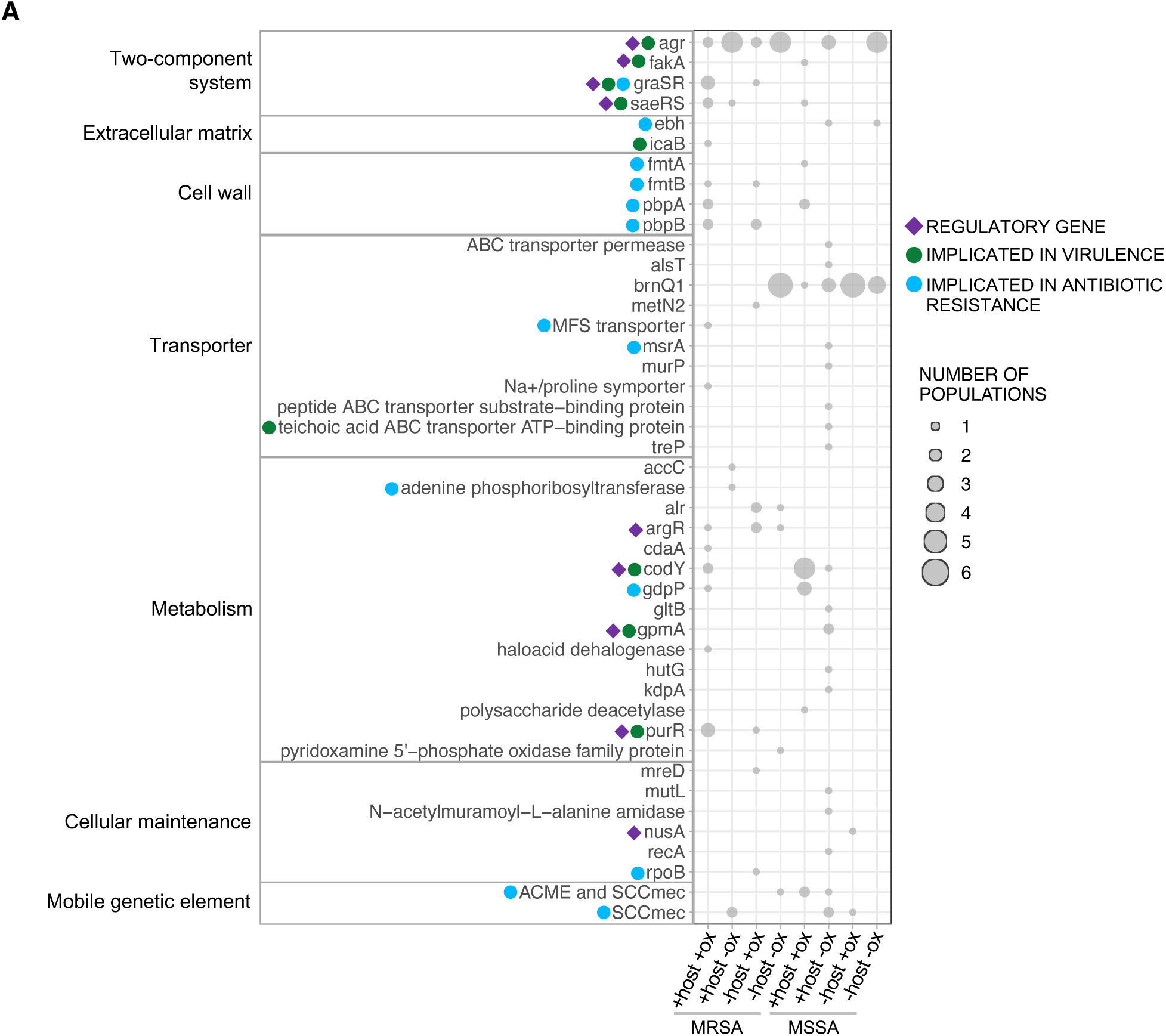
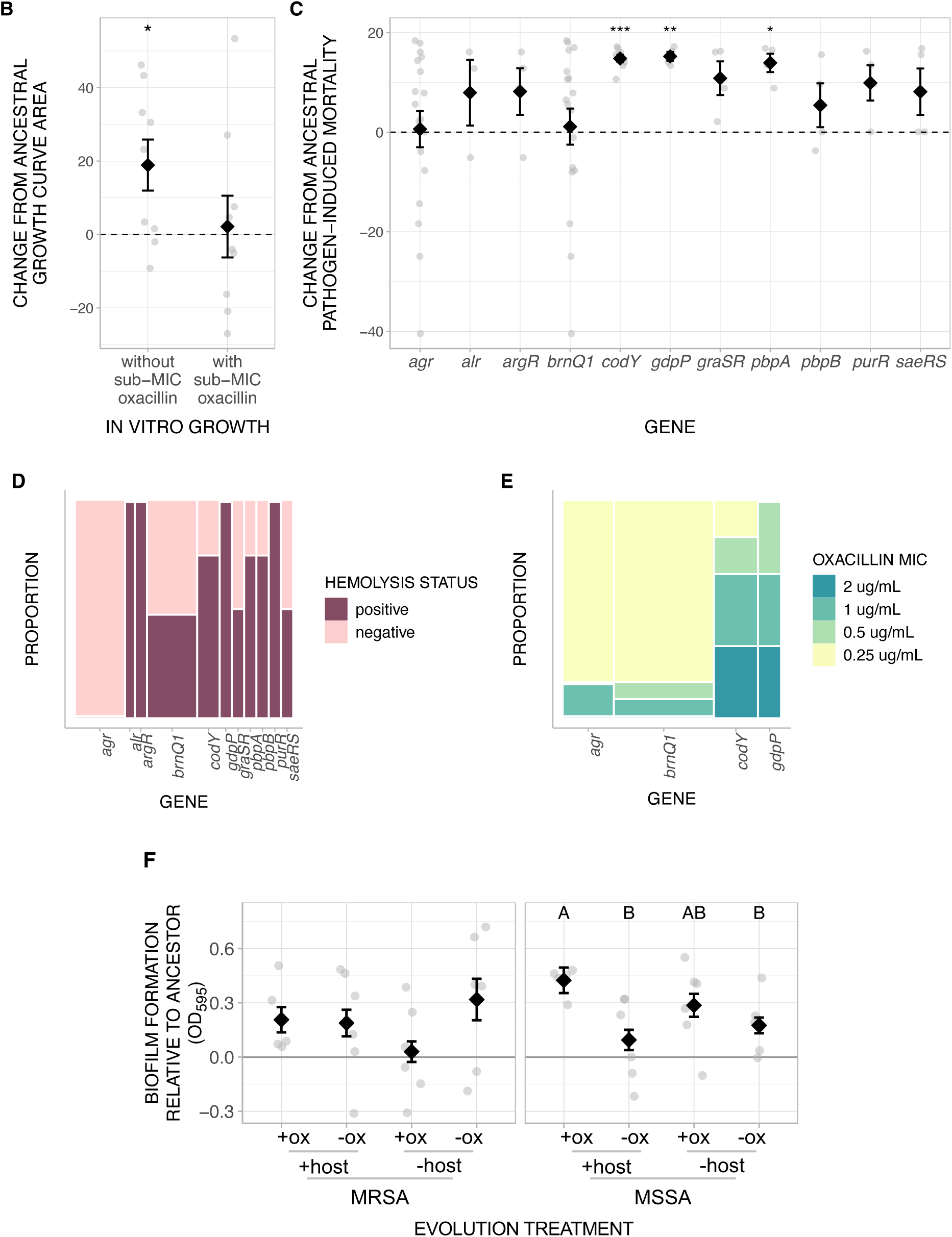
Regulatory genes likely played an important role in pathogen adaptation. A) Mutations swept to fixation, excluding intergenic and synonymous mutations, grouped by general function. The size of each point indicates how many populations had acquired at least one mutation in the gene. Colored shapes next to genes indicate whether these genes are regulatory or have been implicated in virulence or antibiotic resistance in the literature (see Table S1). B) *In vitro* growth of populations with SCC*mec* and ACME deletions with or without sub-MIC oxacillin (one-sample t-test t = 2.73, df = 8, P = 0.026). C) Change from ancestral pathogen-induced mortality *vs*. mutations (*codY* one-sample t-test = 19.56, df = 7, P < 0.001; *gdpP*: 17.02, df = 3, P = 0.002; *pbpA*: 7.54, df = 3, P = 0.012). D) Hemolysis status *vs*. mutations (Fisher’s exact test P < 0.001; *agr vs. alr*: P = 0.036*; argR*: P = 0.006; *codY*: P = 0.005*; gdpP*: P = 0.006*; purR*: P = 0.006). E) Oxacillin MIC *vs*. mutations (Fisher’s exact test P = 0.0015; *codY vs. brnQ1*: P = 0.043). F) Biofilm production of evolved populations. Different letters indicate significant differences (MRSA: 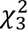 = 7.24, P = 0.06. MSSA: 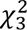 = 17.91, P < 0.001). The column widths in D) and E) corresponds to the number of mutations. Error bars indicate standard errors. All evolved populations were sequenced except for one population from the -host-ox treatment. *P < 0.05, **P < 0.01

Between pathogens with the highest virulence—those under selection from both selective pressures—populations arisen from MRSA had mutations in 25 genes and intergenic regions, while those from MSSA had mutations in 12 genes and intergenic regions. Both treatments had mutations in virulence regulators *codY* (Brinsmade, 2017) and *saeRS* (Montgomery et al., 2010) suggesting some shared pathways to heightened virulence among the two treatments, but mutations in these genes did not occur across all replicate populations. These treatments also had mutations in *gdpP* and *pbpA*, which have been implicated in resistance to beta-lactam antibiotics (Bilyk et al., 2022; Poon et al., 2022), with *gdpP* playing a role in MSSA resistance to oxacillin in particular (Giulieri et al., 2020). Five out of six populations evolved from the MSSA ancestor had a mutation in either gene, which may have contributed to the reduced oxacillin sensitivity in these populations.

Regulatory genes were enriched with mutations (Figure 4A; one-sample Poisson test P < 0.001 for MRSA and MSSA genotypes). These included *agr*, *saeRS,* and *codY*. *agr* is a quorum sensor that regulates the expression of virulence factors (Novick, 2003). All 22 mutations in *agr* were found in populations that have lost the ability to hemolyze red blood cells. In contrast to the other two regulators, *codY* is a transcriptional repressor found in gram-positive bacteria that controls virulence by monitoring nutrient levels in the environment (Brinsmade, 2017).

Nine populations experienced large-scale deletions that included the entire SCC*mec* cassette. Seven of the nine populations were host-associated. Deletion of ACME has been shown to enhance the competitive fitness of *S. aureus* USA300 *in vivo* (Diep et al., 2008). The two MRSA populations that lost their resistance (Figure 3C) had SCC*mec* deletions, suggesting that resistance could be more costly when evolving in a host. The loss of SCC*mec* and ACME was more often identified in populations exhibiting an increase in total growth from the ancestor outside the host (Figure 4B).

### Mutations in antibiotic resistance genes were regularly found in pathogen populations with heightened virulence

We identified mutations arising in specific genes that appeared more often than by chance in measured traits to pinpoint loci potentially underlying evolved phenotypes. We focused on genes where two or more mutations (excluding intergenic or synonymous mutations) had fixed during the experiment regardless of treatment (Figure 4C). Mutations in three genes were regularly identified in populations exhibiting significant increases in virulence from the ancestor: *codY*, *gdpP*, and *pbpA*. Mutations in *agr* in general were not associated with changes in overall virulence, but MSSA populations harboring mutations in this gene were more likely to exhibit greater virulence compared to MRSA populations (Wilcoxon rank sum exact test P = 0.045).

Mutations in specific genes were often found in populations able to hemolyze red blood cells (Figure 4D). There was a greater proportion of populations with *agr* mutations unable to hemolyze red blood cells compared to *alr, argR, codY, gdpP,* and *purR*. There were also significant differences between the mutations regularly identified in oxacillin-resistant populations evolved from the MSSA ancestor (Figure 4E), where a greater proportion of populations with mutations in *codY* had reduced oxacillin susceptibility compared to *brnQ1*. Exposure to oxacillin maintained hemolysis in evolved populations (Figure 2B), suggesting low concentrations of oxacillin exerted negative selection on the *agr* locus. By contrast, mutations in *agr* were often in populations exhibiting loss of hemolytic activity, consistent with previous findings (Schwan et al., 2003).

Because mutations in *codY* appeared to be important for both virulence (Figure 4C) and antibiotic resistance (Figure 4E), we hypothesized that traits controlled by *codY* were responsible for the traits observed in MSSA populations under selection from both pressures. A potential mechanism underlying changes in virulence and antibiotic sensitivity in MSSA may involve biofilm formation. Biofilm is implicated in *S. aureus* pathogenesis as well as in dampening the efficacy of antibiotic treatment (Long et al., 2023). In *C. elegans*, *S. aureus* biofilm enhances virulence and protects the pathogen from host innate immune defenses (Begun et al., 2007). Both sub-MIC level of beta-lactam antibiotics and null-mutants of *codY* induce robust biofilm formation and hemolysis activity (Chen et al., 2021; Kuroda et al., 2007; Majerczyk et al., 2008). Taken together, exposure to sub-MIC level of oxacillin and acquisition of potentially deleterious mutations in *codY* may underlie both increased virulence and reduced antibiotic susceptibility. Our findings partially supported this hypothesis: biofilm formation did not significantly differ for the MRSA genotype. By contrast, biofilm production differed between treatments for the MSSA genotype (Figure 4F), with populations exposed to both selective pressures forming more biofilm than those evolved without host or either pressure, but not those evolved in solely sub-MIC oxacillin. While host and sub-MIC oxacillin selection favored robust biofilms in MSSA, oxacillin by itself also increased biofilm formation, suggesting low antibiotic concentrations contributed significantly to the evolution of drug-sensitive populations. The loci underlying biofilm formation may be different in these two treatments, since mutations in *codY* did not appear in populations under solely oxacillin selection (biofilm is a polygenic trait in *S. aureus* (Zapotoczna et al., 2016)).

### Mutations that arose during experimental evolution are regularly found in strains associated with human systemic infections

We then determined whether mutations that arose in our experiment exist in natural *S. aureus* isolates. We conducted BLAST searches against a dataset of 83,383 high quality public *S. aureus* whole genome assemblies (Raghuram et al., 2024) (Figure 5A) and found matches against the majority of our mutations. We hypothesized that these mutations may be important for *S. aureus* adaptation to different environments. Indeed, for many genes implemented in virulence and antibiotic resistance, a greater proportion of natural isolates containing our mutations were found in blood and systemic infections compared to skin/nose/throat colonization than expected (Figure 5, statistics shown in Table S3). Mutations arisen in MRSA populations under selection from both host and sub-MIC oxacillin had the most matches to natural isolates (Figure 5B), further highlighting the importance of the interaction between these two selective forces. Overall, our results demonstrate that evolution experiments in an invertebrate model can give rise to clinically-relevant mutations in a major bacterial pathogen.

**Figure 5.**
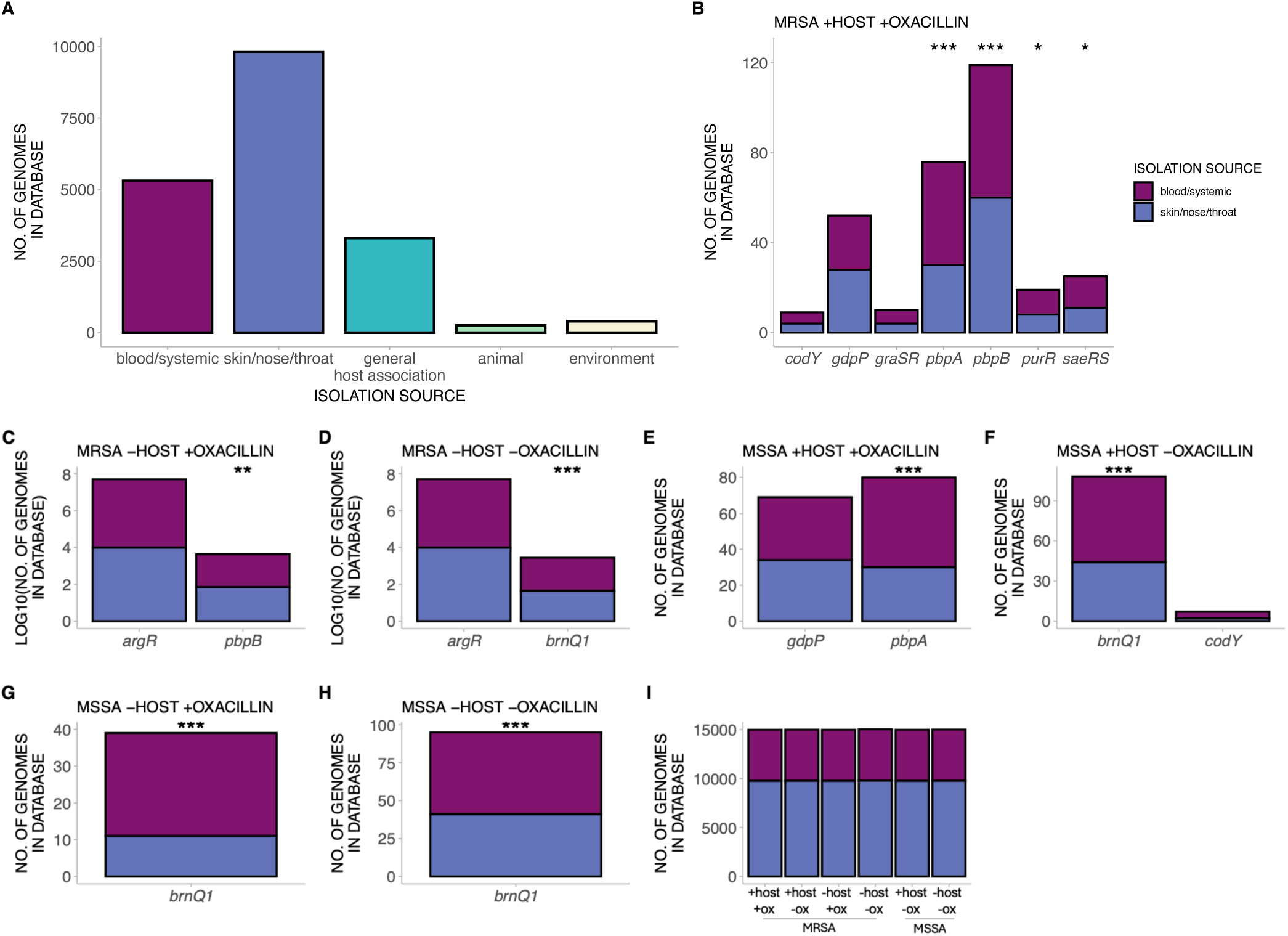
Mutations that arose during experiment are enriched in isolates associated with systemic infection in humans. A) Number of genomes in our public *S. aureus* genome dataset (see Methods) grouped by isolation source. General host-association refers to descriptions not specific enough to assign to other categories. We excluded samples that were ambiguously specified or missing information. B) to H) Number of genomes in the database containing mutations in the respective gene. I) Number of genomes in the dataset containing the mutations arisen in *agr*. This gene is separate from the others to facilitate ease of visualization due to the magnitude of the y-axis. Asterisks indicate significant difference in the proportion of blood/systemic-associated genomes compared to the expected distribution in the dataset. *P < 0.05, **P < 0.01, ***P < 0.001

### MRSA and MSSA exhibited divergent phenotypic and genomic evolution during adaptation

We conducted principal component analysis on the evolved traits (virulence, growth in media with or without sub-MIC oxacillin, and biofilm production) in MRSA and MSSA populations to determine the impact of selective pressures on overall pathogen phenotypic evolution (Figures 6A and 6B). We found a significant effect of treatment for the MRSA genotype, where populations exposed to host and sub-MIC oxacillin clustered together, largely separating from all other treatments (P < 0.05 for all pairwise comparisons). The MSSA genotype had a marginally non-significant effect of treatment. Populations evolved from the MRSA ancestor had similar trajectories for growth in sub-MIC oxacillin and virulence, whereas biofilm production and virulence were more correlated for populations evolved from the MSSA ancestor. For MRSA populations, biofilm production and growth without oxacillin also appeared to be positively correlated.

**Figure 6.**
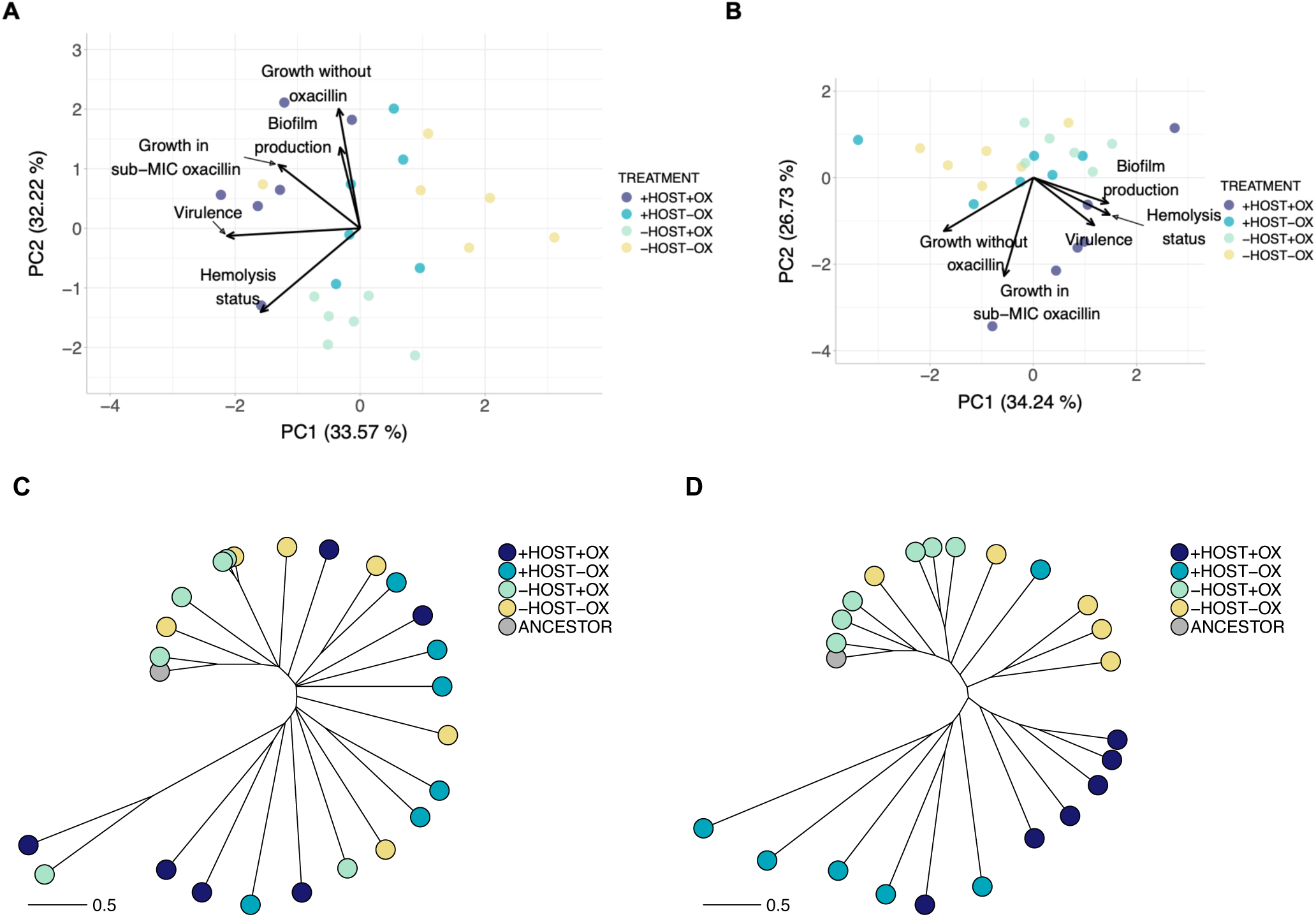
MRSA and MSSA underwent distinct evolutionary trajectories. Principal component analysis of traits evolved from A) MRSA ancestors (PERMANOVA F_(3,23)_ = 8.38, r^2^ = 0.557, P = 0.001) and B) MSSA ancestors (PERMANOVA F_(3,23)_ = 2.20, r^2^ = 0.248, P = 0.064). Phylogenetic tree constructed from frequencies of mutations in populations evolving from C) MRSA ancestors (distance to ancestor: F_(3,20)_ = 2.39, P: 0.099; distance between populations: Kruskal-Wallis 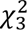 = 21.09, P < 0.001) and D) MSSA ancestors (distance to ancestor: F_(3,19)_ = 10.44, P < 0.001; distance between populations: F_(3,51)_ = 54.20, P < 0.001). Scale bar indicates Euclidean distance.

To determine how host and sub-MIC antibiotic shaped pathogen genomic evolution, we built phylogenies using mutations that arose from the MRSA and MSSA ancestors (Figures 6C and D). We compared Euclidean distances calculated from the frequency of each mutation to determine the genetic distance from the ancestor—how fast each population was evolving—and the genetic distance between populations within each treatment—how much each population had diverged from one another (Betts et al., 2018; Ford et al., 2017). There was no significant difference between treatments for MRSA populations in terms of genetic distance to the ancestor (Figure S6A), suggesting that populations generally exhibited similar evolutionary rates. By contrast, MSSA populations had a significant effect of treatment (Figure S6B), where populations under selection from solely the host had greater genetic distance from the ancestor than populations under selection from solely oxacillin (Tukey’s post-hoc test P < 0.001) or under neither selective pressure (P = 0.004).

For the MSSA group, populations within each treatment tended to exhibit more genetic similarity (Figures 6C and 6D). For MRSA, treatments varied in how divergent populations were (Figure S6C), where populations exposed to both selective pressures diverged more from each other than populations in the other three treatments (Dunn’s post-hoc test P < 0.05 for all pairwise comparisons). Likewise, MSSA populations also differed in how much they varied (Figure S6D). Populations under solely host selection had greater divergence from each other than all other treatments (Tukey’s post-hoc test P < 0.001 for all pairwise comparisons). By contrast, populations under selection solely from sub-MIC oxacillin diverged the least (P < 0.05 for all pairwise comparisons). These findings altogether suggest that even concentrations well below the MIC differentially impacts sensitive and resistant pathogens. As MRSA and MSSA only differed in the presence of an intact *mecA* gene at the start of the experiment, accessory genes may play important roles in shaping bacterial evolution (Jackson et al., 2011).

## DISCUSSION

While antibiotics have existed for millions of years, the mass production and overuse of antibiotics by humans in the last century have exerted an unprecedented level of selection on pathogens (Spagnolo et al., 2021; Wright, 2007). In addition to antibiotics, pathogens face numerous selective pressures such as host defenses and competitors (Bashey, 2015; Cobey, 2014; Kim L Hoang et al., 2024). Opportunistic pathogens are especially likely to encounter diverse selective pressures due to their ability to proliferate outside a host. In this study, we evolved *S. aureus* with or without pre-existing antibiotic resistance under different combinations of host and sub-MIC antibiotics to explore the evolution of virulence and antibiotic resistance. We determined whether increased virulence trades-off with antibiotic resistance by directly selecting for virulence across all host-associated treatments. Nonetheless, we observed differing evolutionary trajectories, where exposure to oxacillin in host-associated treatments resulted in pathogens causing the greatest host mortality. Our experimental design allowed us to tease apart the contributions of the host, sub-MIC antibiotics, and their interaction to demonstrate that adaptation to novel hosts did not require trade-offs in pathogen traits. Indeed, selection from both host and antibiotics actually resulted in rapid pathogen adaptation, potentially creating the conditions under which a pathogen may emerge in new hosts.

Selection from the host and sub-MIC antibiotic favored the greatest virulence and growth in antibiotics, indicating that interactions between these pressures can shape both traits. Sub-MIC antibiotics can facilitate adaptation to high drug concentrations and to the growth environment (Andersson and Hughes, 2014; Pereira et al., 2023). Frequent exposure to low concentrations of antibiotics in *S. aureus* may occur due to its ability to grow and survive in non-host environments (Loeffler et al., 2005; Roberts et al., 2013; Steadmon et al., 2023; Thapaliya et al., 2017), At the cellular level, sub-MIC antibiotics can affect *S. aureus* by directly binding to virulence factors, modulating virulence regulators, and inducing the expression of toxins like hemolysin (Chen et al., 2021; Hodille et al., 2017). Prior exposure to sub-MIC inside and outside hosts can thus mediate traits involved in virulence upon host encounter. Our results further provided evidence for potential links between responses to antibiotic selection and virulence. Mutations in genes involved in resistance to antibiotics were found more often in populations with increased virulence, suggesting that antibiotic adaptation may also favor evolution of virulence. Furthermore, in the absence of oxacillin selection, populations lost their ability to hemolyze red blood cells, supporting previous findings that antibiotics can influence expression of virulence factors (Andersson and Hughes, 2014; Chen et al., 2021).

Likewise, host factors contributed to antibiotic resistance. All populations initially sensitive to oxacillin exhibited increased MICs when under selection from both host and sub-MIC oxacillin. One population evolving under selection from either solely sub-MIC antibiotics or solely the host exhibited decreased oxacillin sensitivity, which is consistent with previous studies examining hosts (McVicker et al., 2014) and sub-MIC (Pereira et al., 2023; Wistrand-Yuen et al., 2018) exposure separately. The pathogens in these studies evolved for much longer than our study, suggesting that having a host accelerated the evolution of antibiotic resistance, potentially due to bottlenecks that favor resistant isolates (McVicker et al., 2014). Evolution in hosts also contributed to pathogen growth outside hosts, an important trait for opportunistic pathogens as persistence in the environment increases the likelihood of contact with new hosts. Host association also resulted in more indels, including large-scale deletions of antibiotic resistance-conferring SCC*mec* and ACME. Genome degradation is a common pattern found in the transition to long-term within-host adaptation across diverse pathogens (Bentley and Parkhill, 2015; Klemm et al., 2016; Lawrence, 2005). Ultimately, exposure to hosts and antibiotics together could create a feedback loop where both evolutionary pressures select for better adaptation in the other, resulting in highly virulent and resistant pathogens that can persist outside hosts.

Several mutations arose in genes with primary roles in metabolism, such as *codY*. It is becoming clear that pathogen metabolism plays a central role in its virulence (Feinbaum et al., 2012; Lindsay et al., 2023) and antibiotic resistance (Zampieri et al., 2017). For example, mutations in chromosomal- and plasmid-encoded metabolic genes are widely prevalent among *E. coli* strains and protect them against antibiotic challenges (Lopatkin et al., 2021; Palomino et al., 2023). Furthermore, antibiotics can directly select for reduced metabolic rates to dampen the deleterious effects of drugs (Lopatkin et al., 2021), and increased resistance is more likely to evolve at high nutrient concentrations (Windels et al., 2024). Similarly, metabolic efficiency can alter virulence (Lindsay et al., 2023), while resource limitations can hinder the evolution of virulence (Pak et al., 2024). In *S. aureus*, CodY de-represses expression of virulence factors under resource-depleted conditions in order to acquire nutrients through the lysing of host cells (Brinsmade, 2017). Our genomic data were consistent with the observation that *codY* may be mediating virulence through control of *agr*: MSSA populations under selection from both host and sub-MIC oxacillin exhibited the greatest virulence and was one of the two treatments without mutations in *agr*. These findings suggested that the *codY* mutations in these populations may be deleterious, rendering CodY less efficient in repressing *agr*-mediated virulence as a result. Future experiments may include introducing these mutations into the ancestral background to directly link the mutations in these genes to evolved virulence. As resources likely differ in abundance and type between hosts and media, our findings illustrated the importance of taking into consideration the environment in which pathogens evolve when evaluating traits of interest.

Regulatory genes in general acquired more mutations than expected by chance alone. This finding supported previous work showing that regulatory elements were more likely to acquire phenotype-altering mutations, especially in negative regulators (e.g., *codY*) (Lind et al., 2015). Enrichment of mutations in regulatory genes has also been found in *E. coli* under antibiotic selection and in *Pseudomonas aeruginosa* under host selection (Card et al., 2021; Jansen et al., 2015). Pleiotropic genes like regulators may increase the capacity to adapt to multiple environments. Because *C. elegans* is a novel host to *S. aureus*, selection is more likely to act on genes that have large initial gains in the fitness landscape, such as those that have pleiotropic effects (Bomblies and Peichel, 2022). Indeed, mutations in several metabolic regulators (*codY, purR, gpmA*) and virulence regulators (*saeRS*) were mainly identified in populations involving hosts. The resulting phenotypes of these mutations, such as increased biofilm production, can then confer multiple benefits like increased virulence and antibiotic resistance. Our experimental results revealed similar patterns to a comparative study analyzing over 2000 *S. aureus* genomes, where mutations in genes with pleiotropic effects, such as *purR*, may have underly pathogen adaptation during systemic infection (Giulieri et al., 2022). Thus, changes in genes of large effects like global regulators appear to be critical mediators between different traits that altogether allow bacteria to rapidly adapt to new and stressful environments.

Antibiotic resistance did not impede the evolution of virulence. Antibiotic resistance tends to be costly in the absence of antibiotics; this cost can subsequently constrain evolution of traits requiring cellular resources, such as virulence (Ferenci, 2016). However, mutations conferring resistance can have little to no costs, depending on the pathogen species and drugs, and costs can be offset by compensatory mutations (Melnyk et al., 2015). Sub-MIC levels favor low-cost mutations (Westhoff et al., 2017), therefore virulence and antibiotic resistance may not trade-off during disease emergence (Visher et al., 2021). Virulence may alternatively be costly in the presence of antibiotic resistance—populations evolved from the MRSA ancestor without either selective pressure exhibited the lowest virulence. The corresponding MSSA populations did not have a similar decline in virulence, suggesting that relaxed selection attenuated virulence for populations with existing resistance. Genomic differences between evolved MRSA and MSSA further demonstrate diverging evolutionary pathways in antibiotic-resistant *versus* antibiotic-sensitive populations. Oxacillin-exposed MSSA had no mutations in *agr* and retained their hemolytic ability, while some MRSA populations gained *agr* mutations and lost the trait, suggesting that other pathways are utilized to express virulence. Lastly, mutations in MRSA under selection from both host and antibiotics were significantly enriched in blood and systemic *S. aureus* isolated from humans, suggesting that antibiotics may have significant roles in MRSA invasive infection. Acquiring mutations in genes that impact antibiotic resistance, such as *pbpA* and *pbpB*, may allow *S. aureus* to persist longer in hosts. Additional direct tests are needed to evaluate the role of these mutations in adaptation of *S. aureus* to different infection sites.

Recent work has shown that diverse genotypes of one pathogen species converge on similar phenotypes when facing similar selective pressures (Filipow et al., 2023). Here, we showed that antibiotic resistance status can lead otherwise isogenic pathogens to use different adaptive strategies. Initially susceptible pathogens, facing novel selection from host and antibiotics, exhibited reduced phenotypic and genomic variation over evolutionary time and thus may have had a limited number of pathways to adapt to new environments. By contrast, resistant populations had greater variance and thus may have had more routes to adapt. Overall, the potential for low-cost (Visher et al., 2021; Westhoff et al., 2017) and pleiotropic mutations (Bomblies and Peichel, 2022) may have created ideal conditions for a novel pathogen to establish in host populations regardless of its susceptibility to antibiotics. Our work demonstrated that responding to multiple pressures can result in rapid adaptation to new hosts and even evolution of traits not under selection, emphasizing the importance of non-canonical pathways (e.g., metabolism) in shaping these traits. Pathogen evolution in a tractable invertebrate animal model yielded phenotypes and genotypes similar to those identified in mammalian hosts, highlighting the utility of evolution experiments to identify potential ecological and genetic mechanisms that may give rise to pathogen traits conserved across systems. Our findings ultimately emphasize the importance of considering the host context in the evolution of antibiotic resistance. Integrating multiple traits, such as virulence, antibiotic resistance, and fitness may be critical in identifying the factors that facilitate host shifts and persistence of drug-resistant pathogens.

## ACKNOWLEDGEMENTS

The authors are grateful for insightful discussion with members of the Read and Morran labs, and Shaun Brinsmade for reading the manuscript. We thank Carmen Alvarez, Jason Chen, and Emily Stevens for help with experimental protocols.

## AUTHOR CONTRIBUTIONS

MS, LTM, and TDR conceptualized the project. MS, KLH, MP, MHD, and JDG conducted the experiments. KLH and MS analyzed the data and drafted the manuscript. All authors contributed to manuscript revision.

## DATA AVAILABILITY

Raw sequences have been deposited in the NCBI Sequence Read Archive under the BioProject accession number PRJNA1162848, and phenotypic data deposited in the Figshare Repository (10.6084/m9.figshare.28745558).

## MATERIALS AND METHODS

### Bacteria and nematode strains

*Staphylococcus aureus* USA300 JE2 (NR-46543) and USA300 JE2 *mecA*-tn (Transposon Mutant NE1868) from the Nebraska Transposon Mutant Library were acquired from BEI resources (https://www.beiresources.org/). To ensure true isogenic ancestors, the *mecA*-tn was transduced into the USA300 JE2 background, selecting for the erythromycin resistance marker on the Mariner transposon, and the insertion site was confirmed by polymerase chain reaction. Transposon maintenance was confirmed by testing for resistance to erythromycin (5µg/mL) or by genome sequencing when available. In cases where transposon loss was verified by genome sequencing, strains were tested and shown to maintain sensitivity to oxacillin. As the primary function of the transposon was to knock out *mecA* function, loss of the transposon should not confound subsequent analyses as antibiotic sensitivity was maintained.

*Caenorhabditis elegans* strain N2 Bristol and *Escherichia coli* strain OP50 were provided by the Caenorhabditis Genetics Center, which is funded by the NIH Office of Research Infrastructure Programs (P40 OD010440). Nematodes were maintained on Nematode Growth Medium Lite (US Biological, Swampscott, MA) according to WormBook protocols (Stiernagle, 2006).

### Experimental evolution

Selection for increased virulence of *S. aureus* (USA300 JE2 or USA300 JE2 *mecA-tn*) in *C. elegans* was performed by passaging *S. aureus* from dead hosts 24 hours post-infection. Half of the *S. aureus* populations were additionally subjected to antibiotic pressure during the passages before and after *C. elegans* selection with sub-MIC oxacillin exposure (0.03125µg/mL). This concentration was tested against both ancestral strains and did not substantially inhibit growth of the methicillin-sensitive USA300 JE2 *mecA*-tn strain.

For each passage of experimental evolution, *S. aureus* was grown in Brain Heart Infusion (BHI) broth with 4µg/mL colistin (to prevent *E. coli* contamination from the nematode intestine) ± 0.03125µg/mL oxacillin and incubated at 37°C overnight, shaking at 250rpm. The next day, 200µL of the overnight *S. aureus* culture was plated onto BHI agar and incubated at 28°C overnight. Approximately 1500 *C. elegans* were plated and allowed to consume *S. aureus* for 24 hours before 30 dead nematodes were picked from each plate. Nematodes were identified as dead by a lack of response to tapping by a platinum wire (Ford et al., 2017; Kim L Hoang et al., 2024). Bacterial extraction methods were adapted from (Vega and Gore, 2017). In brief, picked dead nematodes were washed once with M9 buffer supplemented with 0.1% Triton X-100, once with M9 buffer, then subjected to a 1:1000 bleach-M9 buffer solution for 15 minutes. Nematodes were then washed twice with M9 buffer and ground with a motorized pestle to extract *S. aureus* from the host intestine. *S aureus* populations evolving without hosts were obtained from nematode-free bacterial lawns with a loop. Populations of *S. aureus* were grown as previously before freezing and storage at −80°C. Twenty-five percent of each frozen culture was used as inoculum for the next round of experimental evolution. Preliminary experiments determined colony-forming units per milliliter dropped by approximately 50% after freezing. To maintain population diversity, the inoculum amount (25%), adjusted for freezing, was chosen to be within the dilution ratio presented in (Wahl et al., 2002) that minimizes the chance that rare beneficial mutations are lost. Twelve rounds of passage were completed for 2 *S. aureus* genotypes x 2 host treatments x 2 antibiotic treatments x 6 replicates = 48 populations. Whole populations were frozen for subsequent analyses. We conducted all assays to evaluate changes in evolved *S. aureus* at the population level instead of single isolates to represent the context under which *S. aureus* evolved in our experiment and generally how pathogens exist in nature and in hosts (Cordero et al., 2012; Launay et al., 2021; McAdam et al., 2011; Mei et al., 2021; Paterson et al., 2015). Population level assays also allow for potential interactions between genotypes that give rise to phenotypes like virulence (Ruiz-Bedoya et al., 2023) and antibiotic resistance (Azimi et al., 2020; Cordero et al., 2012) that would otherwise not be captured in single-isolate assays.

### Mortality assay

Nematodes were age-synchronized as in (Penley and Morran, 2018) and grown on *E.coli* OP50 until L4 larval stage (48 hours at 20°C). Nematodes were subsequently washed off, counted, and approximately 200 individuals were plated on *S. aureus* lawns on BHI plates. Another set of nematodes were plated on *E. coli* OP50 to estimate the total number of nematodes. Plates of *S. aureus* were prepared 24 hours prior by plating 200µL of an overnight population culture on BHI agar and grown at 28°C. After two days at 20°C, plates were scored by counting live nematodes. Host mortality data were analyzed using a generalized linear model with a binomial distribution and followed by Tukey multiple-comparison tests to determine pairwise differences. To compare variances across treatments, we conducted Levene’s test for homogeneity of variance. All statistical analyses, including those below, were conducted in R (version 4.2.0).

### Hemolysis phenotype assay

Populations of *S. aureus* were streaked onto Trypticase Soy Agar II with 5% sheep’s blood plates and incubated overnight at 37°C. Bacteria were assessed as either hemolysis positive or negative depending on whether there was a complete clearing around the colonies. We did not cross-streak with RN4220, which produces beta-hemolysin, and allows for detection of delta-hemolysin. Therefore, our assay mainly detects alpha-toxin and phenol-soluble modulins. Hemolysis data were analyzed using a generalized linear model with a binomial distribution. To determine whether there was an association between hemolysis status and host mortality, we compared the host mortality means of hemolysis-positive vs. hemolysis-negative pathogens using a Kruskal–Wallis test.

To determine whether there was variation in hemolysis ability within populations, we streaked 10 colonies from each of the 48 evolved populations onto Trypticase Soy Agar II with 5% sheep’s blood plates and incubated overnight at 37°C. To compare the number of isolates able to hemolyze, we conducted a generalized linear model with a binomial distribution. To compare variances, we conducted Levene’s test for homogeneity of variance.

### *In vitro* growth assay

We measured growth in BHI to simulate conditions under which pathogens were grown in our experiment. Rich media allowed us to determine growth when conditions are optimal for evolved pathogens. Measuring growth in minimal media, while being more similar to conditions that *S. aureus* may face in its environment, would introduce another selective pressure. Overnight cultures of each evolved population was diluted 1:1000 in BHI broth, then 200µl of each dilution was added to 96-well plates, covered with a Breathe-Easy sealing membrane (Sigma) to minimize evaporation. The optical density at 600nm at 37°C with shaking at 282 cycles per minute was recorded every 10 minutes for 16 hours (1000 minutes) using a BioTek Synergy HTX Multimode Reader. We randomly assigned each of the 48 evolved populations to the inner wells of the 96-well plates, with at least three technical replicates per population. Each plate also included technical replicates of each ancestor. Each ancestral strain was assayed with four technical replicates per plate across four plates. These steps were repeated for media with 0.03125µg/mL oxacillin. We used the R package *gcplyr* (Blazanin, 2023) to calculate the area under the curve of density, a metric that quantifies the overall bacterial growth. We compared the growth of populations across treatments using the mean area under the curve of the technical replicates for each population. We used a linear model followed by Tukey multiple-comparison tests to determine pairwise differences between the ancestors, and linear mixed models for evolved populations grown with or without oxacillin.

### Antimicrobial susceptibility assay

Antimicrobial susceptibility testing for oxacillin was done according to Clinical & Laboratory Standards Institute (CLSI) standard protocols for broth microdilution (CLSI, n.d.). In brief, populations were streaked from frozen stock and re-streaked the next day. From the second day culture, populations were resuspended in normal saline to a 0.5 McFarland standard. Then, these cultures were diluted 1:20 in normal saline before 10µL was added to 90µL of CAMHB + 2% NaCl with the appropriate concentration of oxacillin. Plates were incubated at 35°C without shaking and read at 24 hours. For statistical analysis of populations evolved from the MSSA ancestor, we set a threshold such that anything above the ancestral susceptibility level (0.25ug/ml) was considered a decrease in oxacillin sensitivity. We then compared the proportion of populations exhibiting a decrease in oxacillin sensitivity with those that had not using Fisher’s exact test with false discovery rate-adjusted (FDR) p-values.

To determine whether there was variation in antibiotic susceptibility within evolved populations, we sampled 10 colonies from each of the four populations with the most number of mutations (two from MRSA and two from MSSA), and two additional randomly selected populations. We argued that variation would be the most observable in populations with the most genetic diversity. We then followed the standard protocol for a Kirby-Bauer disk diffusion susceptibility test (Hudzicki, 2009) using oxacillin. Briefly, we inoculated Mueller-Hinton agar plates (Hardy Diagnostics) of overnight cultures with standardized OD_600_ and placed an oxacillin-impregnated disk (HardyDisk AST Oxacillin, OX1) onto each plate. We measured the diameter of the zone of inhibition after incubation at 37°C overnight.

### Biofilm assay

Overnight cultures of each population were diluted 1:40 in BHI broth containing 0.5% glucose, then 100µl of each dilution was added to the inner wells of 96-well plates (NEST Biotechnology, flat-bottom, tissue culture-treated). Plates were incubated for 24 hours at 20°C, the temperature that nematodes experience during infection. Each well was washed three times with 200µl of phosphate-buffered saline (PBS) to remove non-adherent cells. We then stained cells with 125µl of 0.1% crystal violet for 20 minutes, followed by washing each well twice with 200µl PBS. We then resuspended the crystal violet in 125µl of 30% acetic acid. After incubation for 10 minutes, we measured the optical density at 595nm using a BioTek Synergy HTX Multimode Reader. We randomly assigned each of the 48 evolved populations to the inner wells of the 96-well plates, with at least three technical replicates per population. Each plate also included technical replicates of each ancestor. We standardized the mean of technical replicates for each population against the respective ancestor within each plate to control for variation across plates. We used mixed linear models followed by Tukey multiple-comparison tests to assess pairwise differences between treatments.

### Whole-genome shotgun sequencing and variant annotation

Bacterial genome DNA was extracted from whole populations using Wizard Genomic DNA Purification Kit (Promega). Population sequencing allows us to capture the genetic variation within populations, which is critical to understanding how pathogens evolve (Azimi et al., 2020; Ford et al., 2017, 2016; Kim L Hoang et al., 2024; Kim L. Hoang et al., 2024; Mahrt et al., 2021). Genome sequencing of these DNA samples was performed at the M-Gen center using a paired-end library preparation method based on Illumina Nextera and sequenced on the NextSeq 550.

The raw data was quality trimmed and adapters removed using illumina-cleanup (https://github.com/rpetit3/illumina-cleanup) or with bactopia (Petit and Read, 2020). Mutations were called based on comparison to the reference USA300 JE2 genome (BioProject: PRJNA224116, BioSample: SAMN06677988, Assembly: GCF_002085525.1) using breseq (Deatherage and Barrick, 2014). The presence of large-scale deletions and other genetic changes were confirmed by analysis of de novo assembled genomes performed by the SKESA assembler software (Souvorov et al., 2018). The ancestral populations were almost identical to the reference genome used except for three mutations that were not found in any evolved populations. Another three mutations in *bioD1, cls* and *ccdC* were present in all populations derived from the MSSA ancestor and likely occurred in the first passage from the frozen culture before experimental evolution was performed. We therefore removed these six mutations from the analysis. Raw sequences have been deposited in the NCBI Sequence Read Archive under the BioProject accession number PRJNA1162848.

The number of mutations (excluding synonymous or intergenic regions) across treatments were analyzed using a generalized linear model with a Poisson distribution. For mutation enrichment analysis, we followed the steps as described previously (Lees et al., 2017). Briefly, we calculated the “mutation rate” for each ancestor by dividing the total number of mutations by the reference genome size. We multiplied the mutation rate by the total length of regulatory genes (Ibarra et al., 2013) with mutations in our study to generate the expected number of mutations for these genes. We then conducted single-tailed Poisson tests to determine whether the actual number of mutations in regulatory genes was greater than expected.

For phenotypes *vs.* mutations, we considered only mutations occurring at least twice throughout the experiment. For host mortality *vs.* mutations and growth rate *vs.* SCC*mec* and ACME, we conducted a one sample t-test for each evolved pathogen against its respective ancestor. P-values were adjusted using the FDR. For hemolysis status *vs.* mutations and oxacillin MIC *vs.* mutations, we conducted Fisher’s exact tests with Bonferroni corrections.

### Comparing mutations in experiment against publicly available genomes

For each non-synonymous mutation in each gene, we created the FASTA file of the USA300 JE2 protein product and a cognate file of the mutant protein with the amino acid substitution edited manually. We ran tBLASTn (version 2.12.0) searches of both proteins against 83,383 high quality *S. aureus* whole genome assemblies (Raghuram et al., 2024). We filtered out matches to genomes with less than 95% identity. We screened for cases where mutants had a better match to a genome than the ancestral protein (higher bitscore).

We collected meta-data for BioSamples accession linked to each genome from NCBI Entrez using a Python script from Jason Stajich (https://github.com/stajichlab/biosample_metadata), then manually curated the “isolation_source” column to either blood/systemic infection, skin/nose/throat, general host-association, animal, environment, others, and missing (see Table S4 for specific terms). For BioSamples matching mutants within each treatment, we compared the proportion of blood/systemic infection *vs.* skin/nose/throat against the expected proportion (all BioSamples in the dataset) using a chi-square goodness-of-fit test.

### Principal component analysis

We conducted a principal component analysis on the phenotypes quantified in evolved populations using the prcomp function in R for each *S. aureus* genotype. We performed permutational analysis of variances (PERMANOVA, 999 permutations) to test for differences between treatments using adonis2 of the *vegan* package (Oksanen et al., 2024) and pairwiseAdonis (Martinez Arbizu, 2020).

### Genetic distance

We generated a matrix of Euclidian genetic distances (the square root of pairwise differences) using the frequencies of mutations that arose for each *S. aureus* genotype, from which we constructed phylogenies (Betts et al., 2018; Ford et al., 2017). We used pairwise differences between the ancestor and each evolved populations to compare rates of evolution across treatments, and pairwise differences between replicate populations within each treatment to compare the degree of divergence across treatments. We compared rates of evolution for MRSA and MSSA treatments, and degree of divergence for MSSA treatments, using linear models followed by Tukey multiple-comparison tests. We compared the degree of divergence for MRSA treatments using a Kruskal–Wallis test followed by Dunn’s post-hoc test.

## SUPPLEMENTAL FIGURES

**Figure S1.**
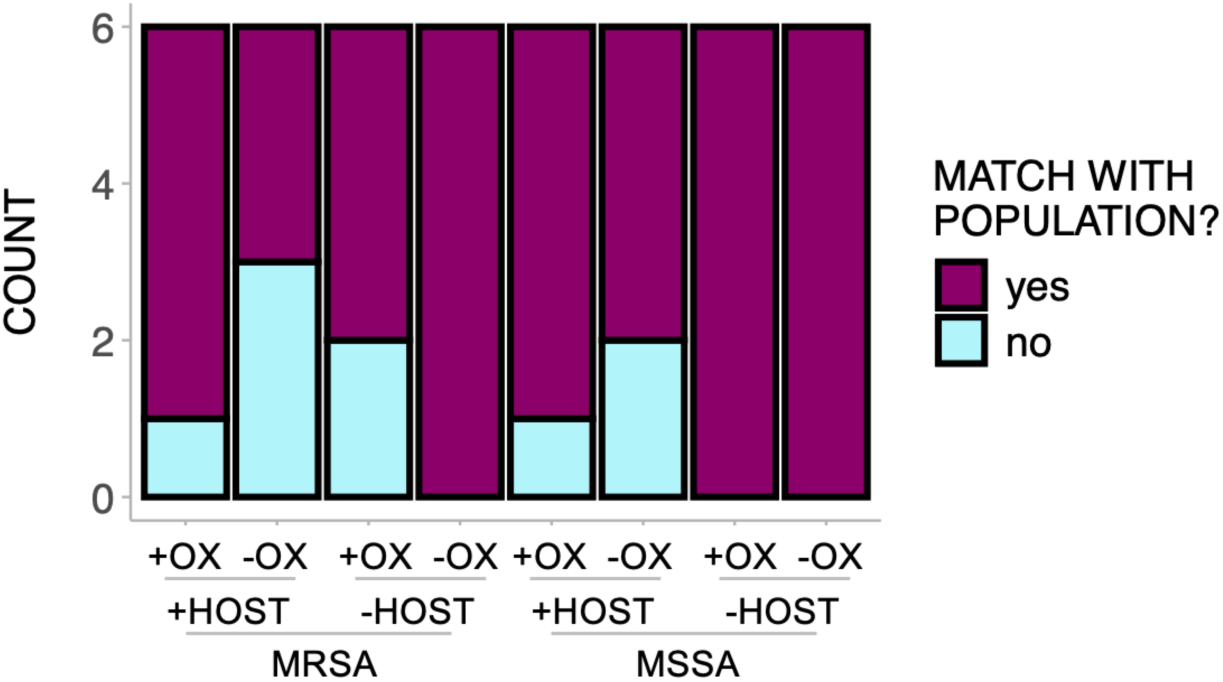
Most colonies sampled matched their respective population in terms of the ability to hemolyze sheep’s blood (*i.e.,* over 50% of colonies having the same hemolysis status as the population they were sampled from).

**Figure S2.**
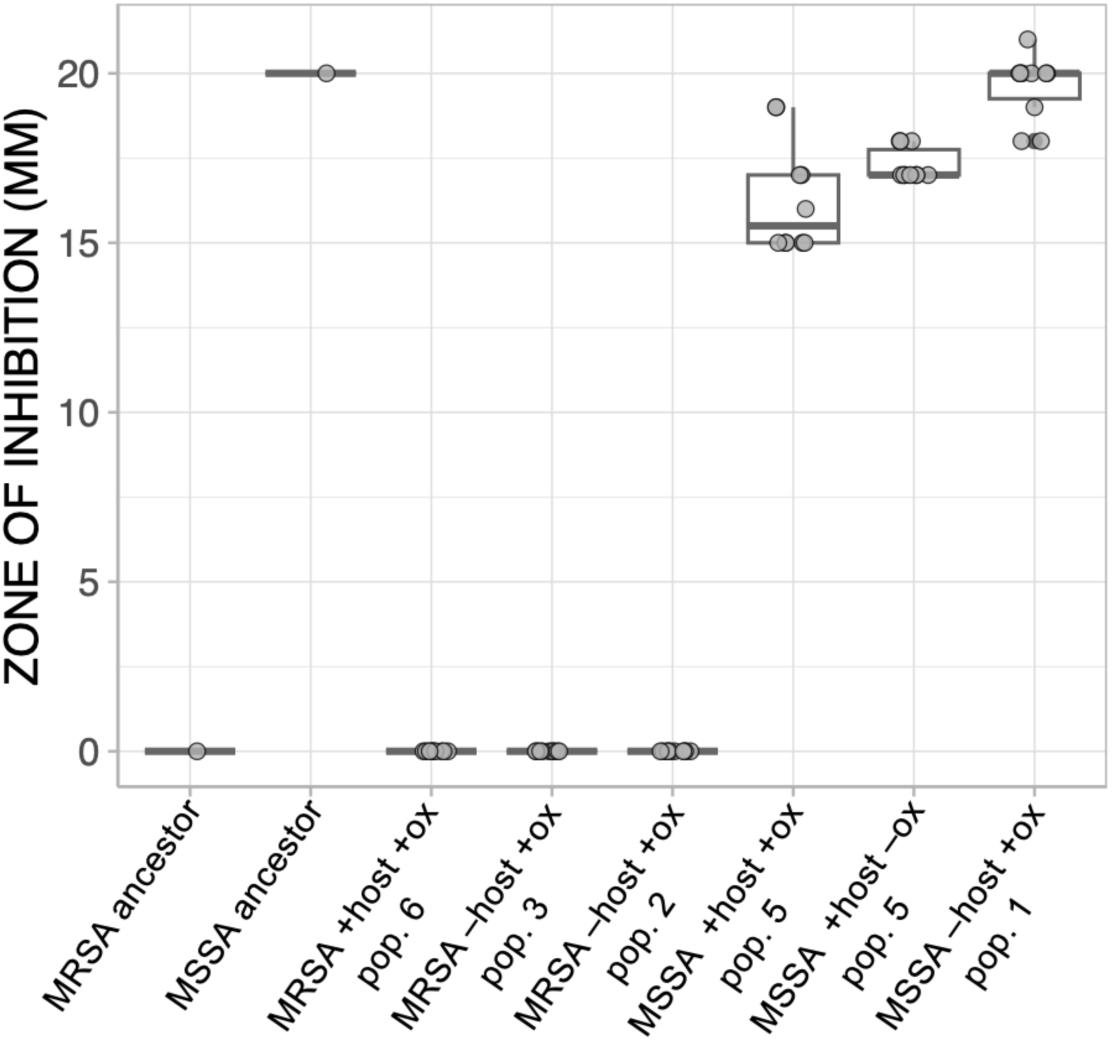
Zone of inhibition from Kirby-Bauer disk diffusion susceptibility test with oxacillin for colonies sampled from each of the four populations with the most number of mutations (two from MRSA and two from MSSA), and two additional randomly selected populations.

**Figure S3.**
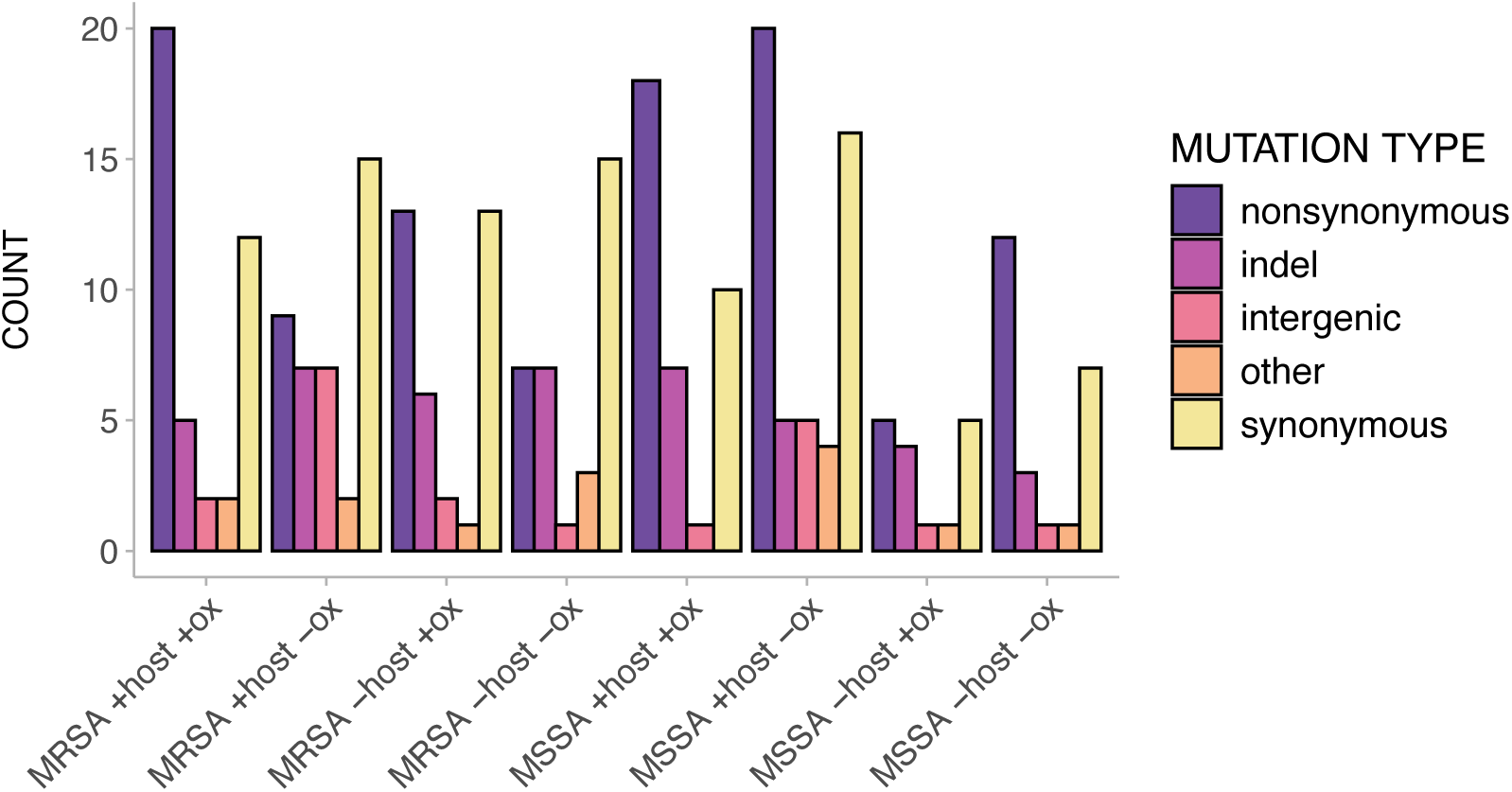
Count of all mutations (from 10-100% frequency) arisen in evolved populations.

**Figure S4.**
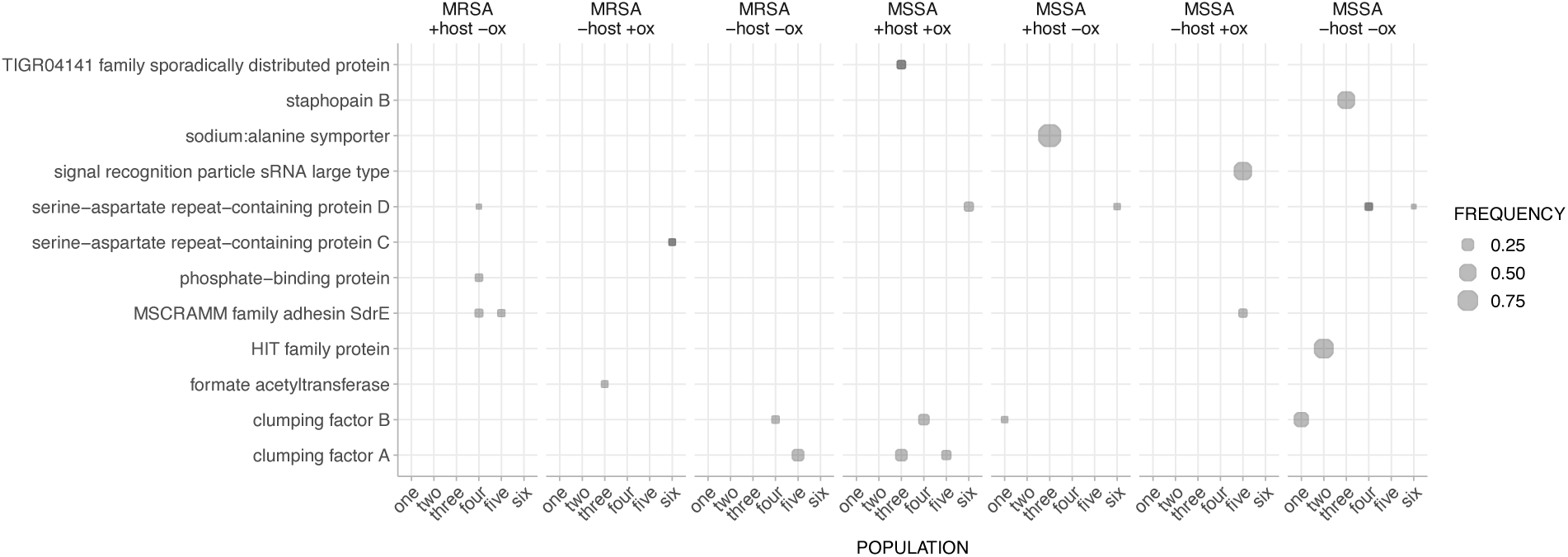
Mutations at frequencies between 0.1 and < 1, excluding synonymous or intergenic mutations, in each evolved population. Points with darker shades indicate more than one mutation present. All mutations in MRSA +host +ox populations between 0.1 and < 1 are all either synonymous or intergenic. A table of all mutations and frequencies is available on figshare (10.6084/m9.figshare.28745558)

**Figure S5.**
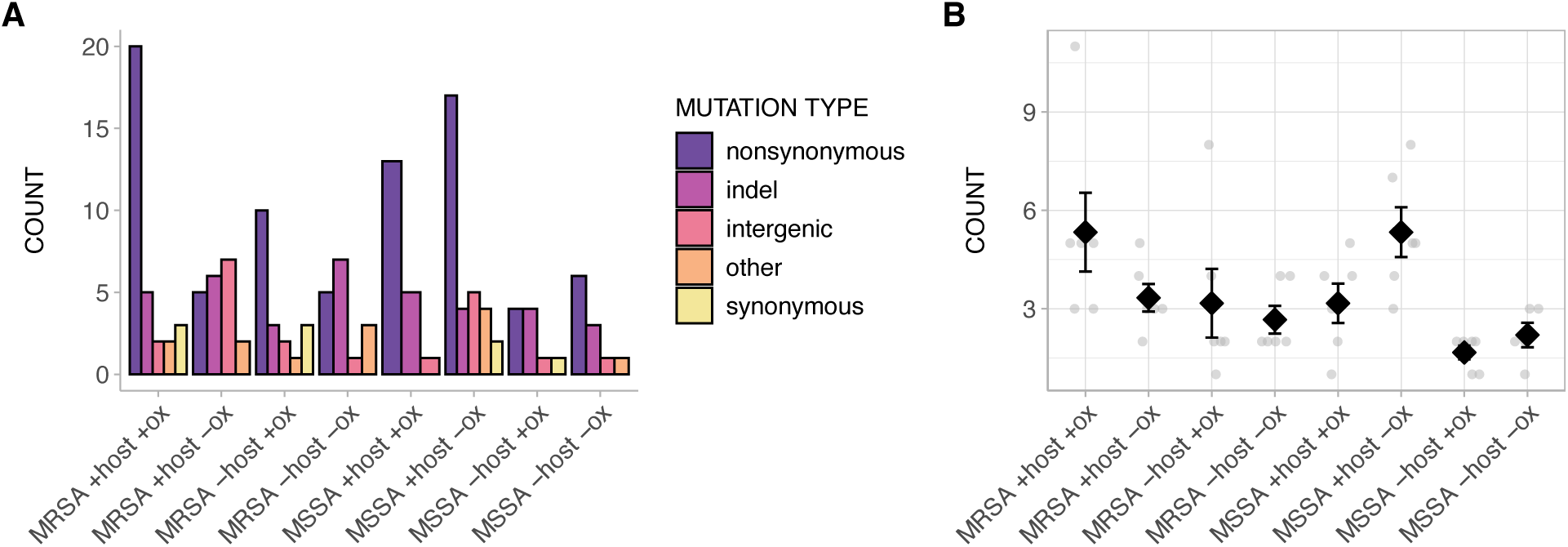
A) Count and B) Mean of mutations swept to fixation in evolved populations.

**Figure S6.**
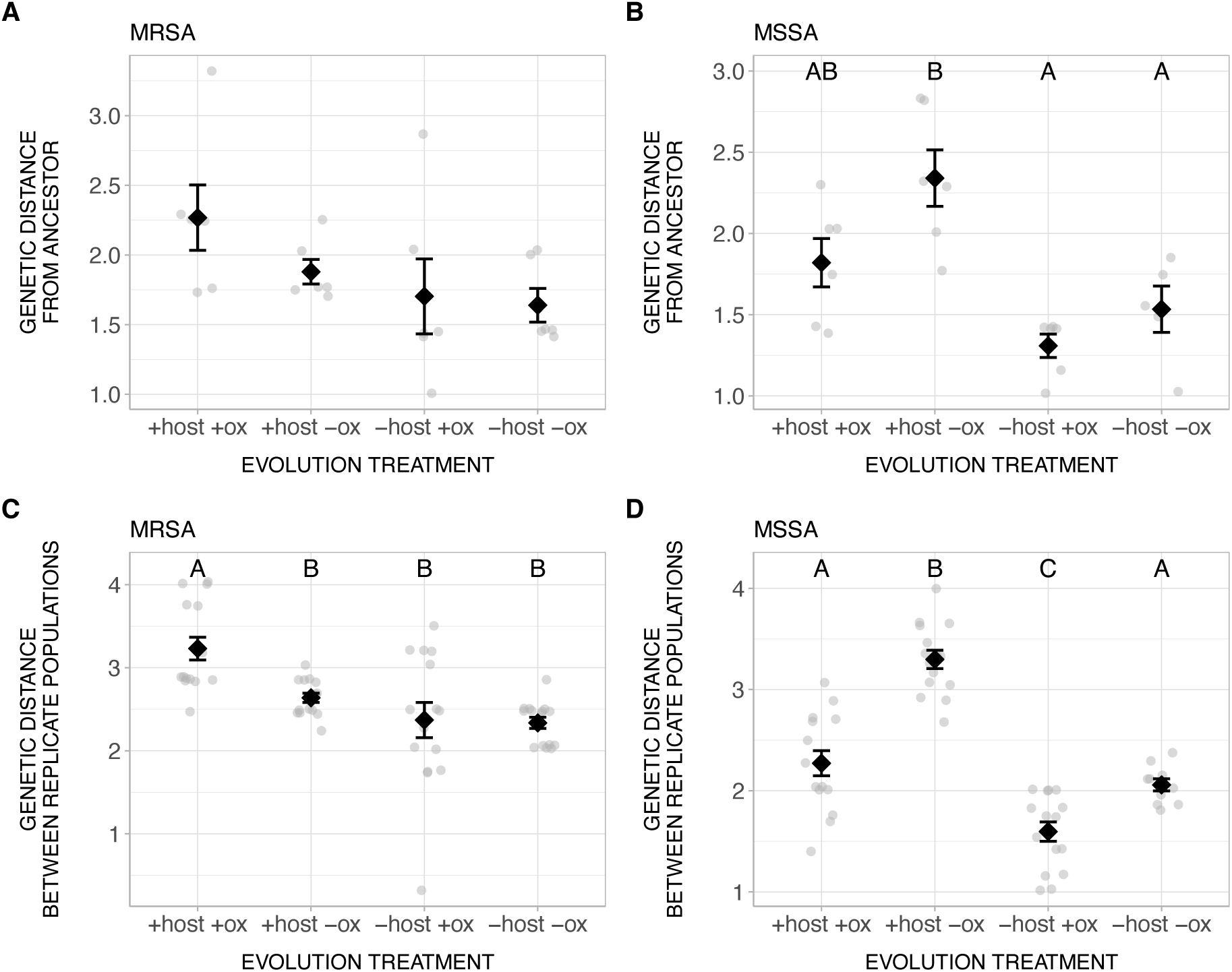
Genetic distance between the ancestor and evolved populations for A) MRSA and B) MSSA genotypes. Genetic distance between replicate populations within each treatment for C) MRSA and D) MSSA genotypes. Different letters indicate significant differences. Error bars indicate standard errors.

## SUPPLEMENTAL TABLES

**Table S1.**
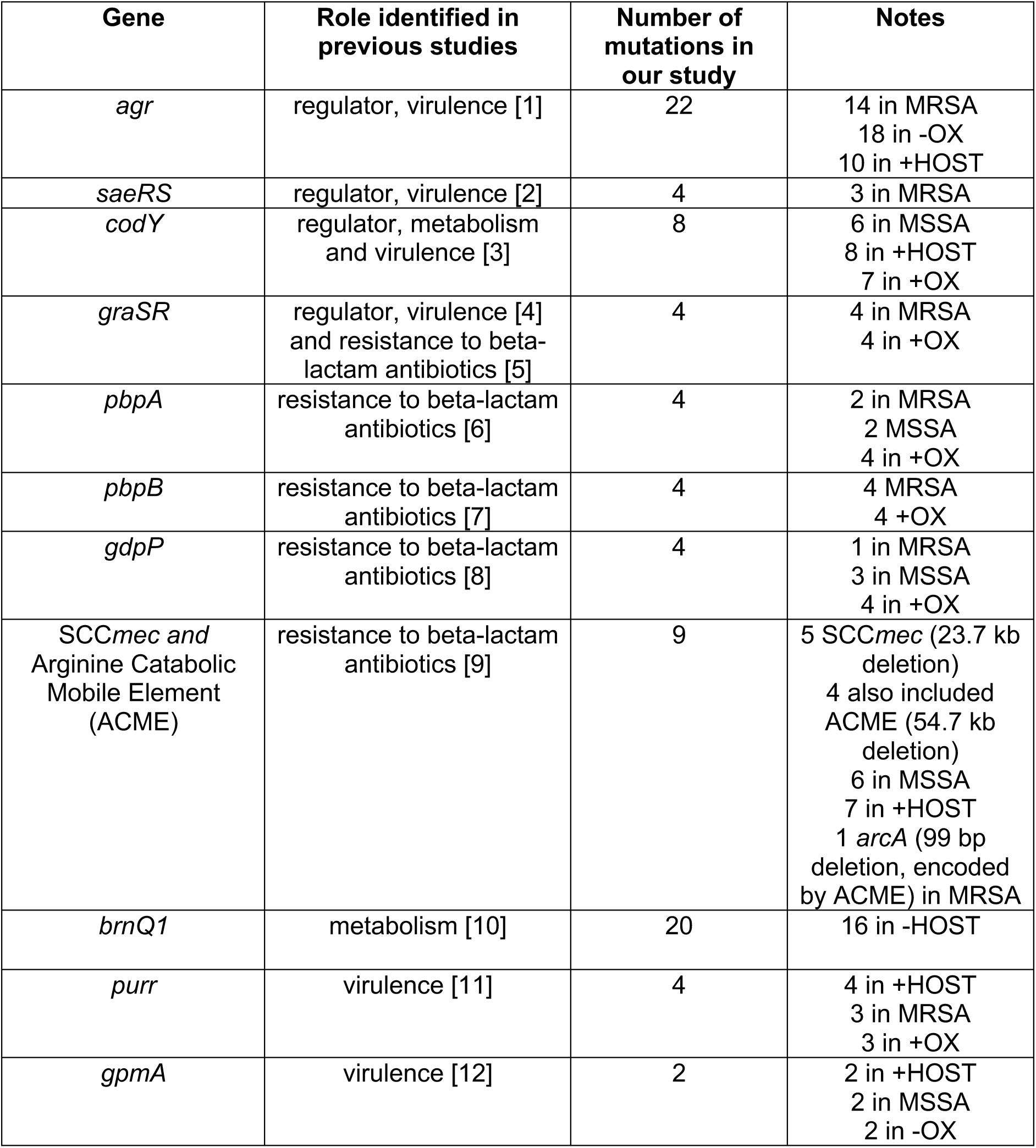
Nonsynonymous mutations occurred in genes with known roles in virulence and antibiotic resistance.

**Table S2.**
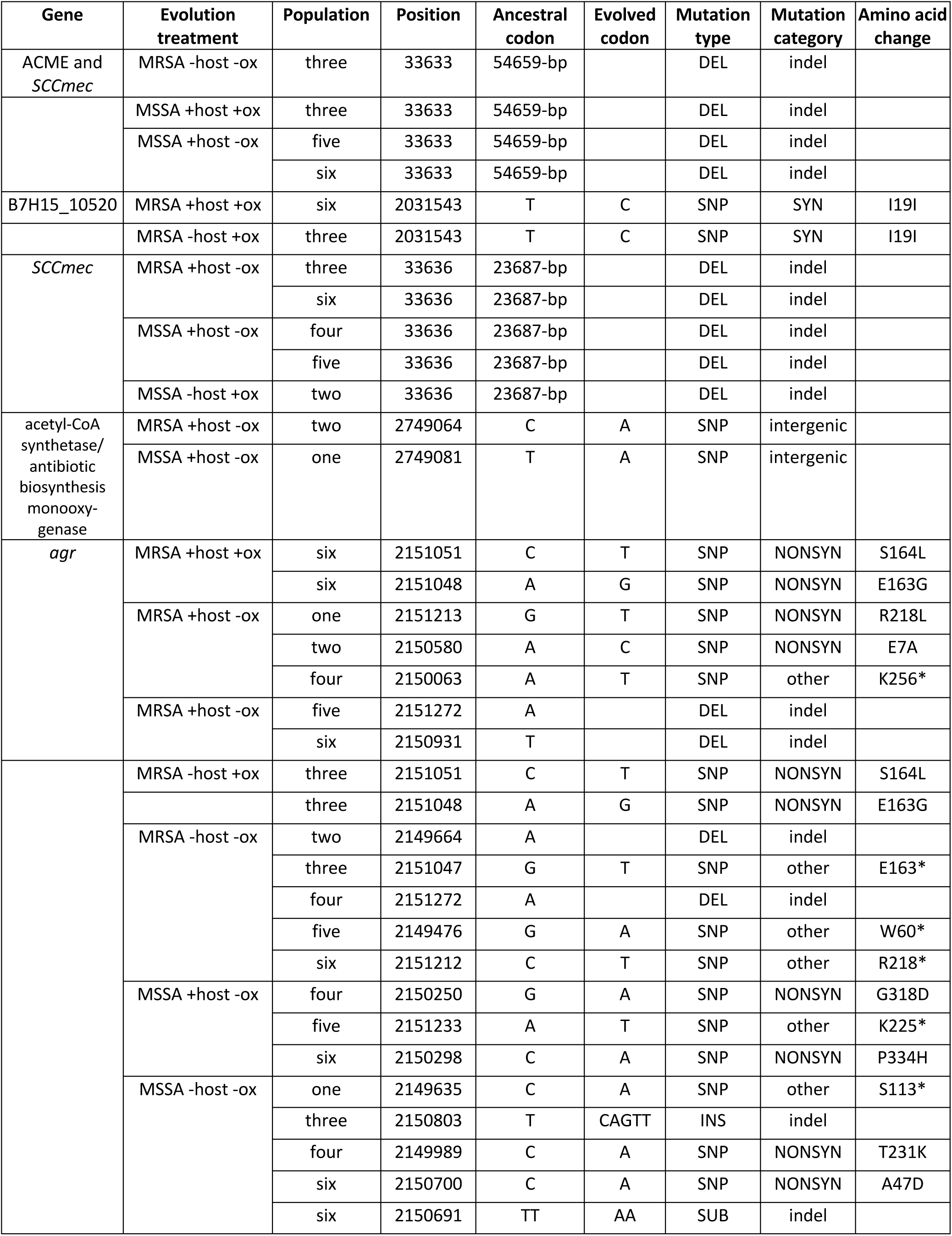
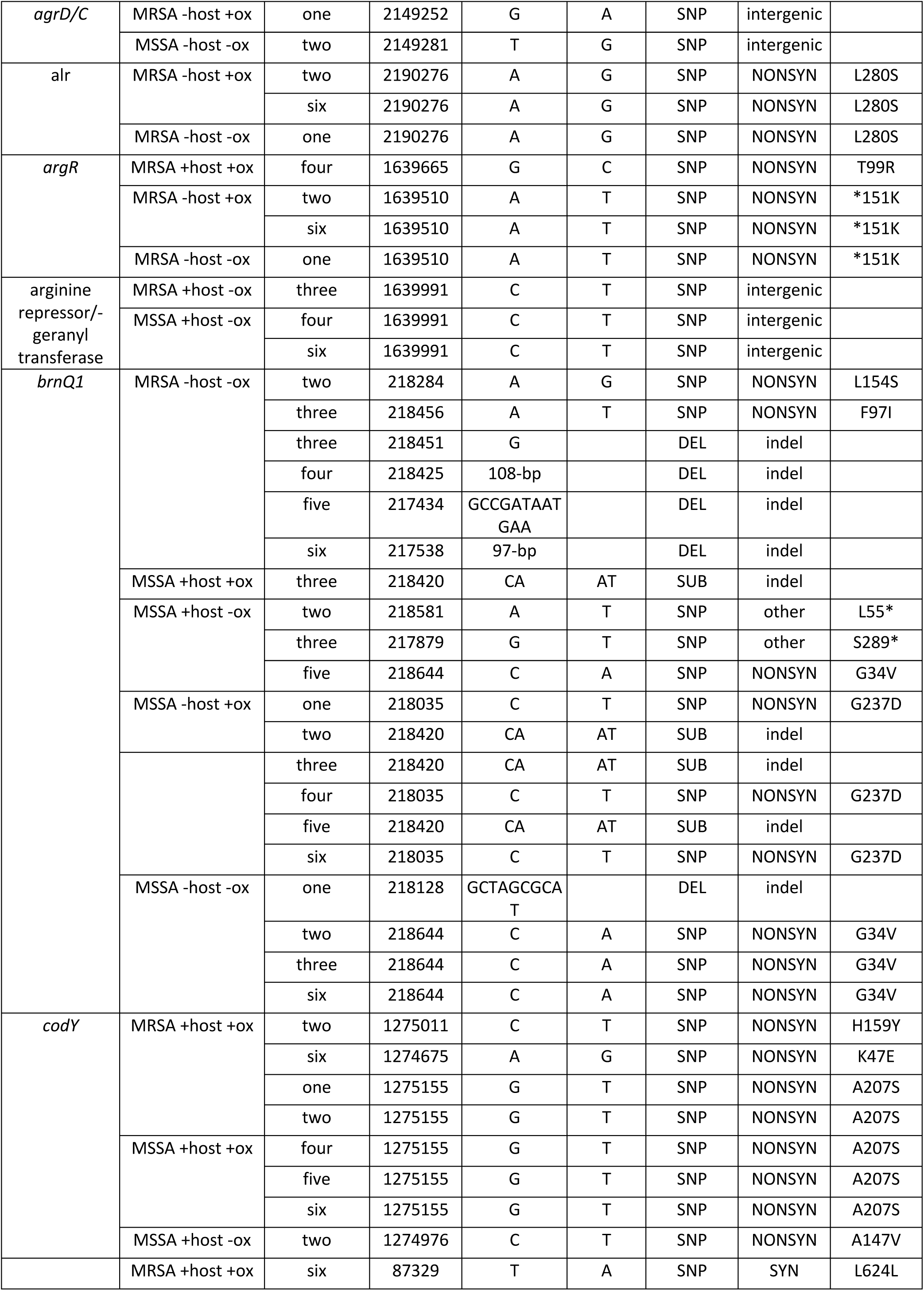
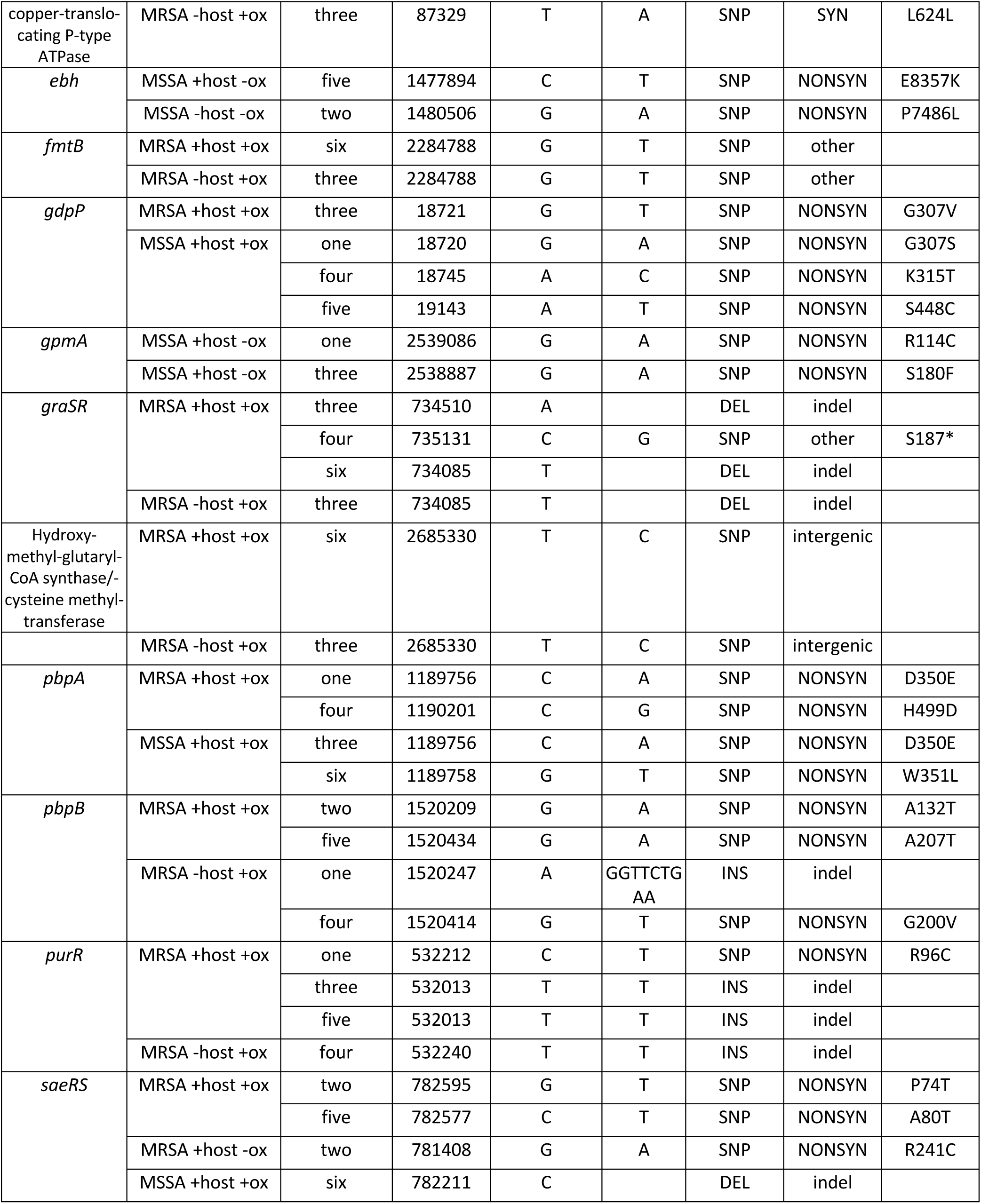
Fixed mutations in genes and intergenic regions appearing in more than two populations. SYN = synonymous, NONSYN = nonsynonymous.

**Table S3.**
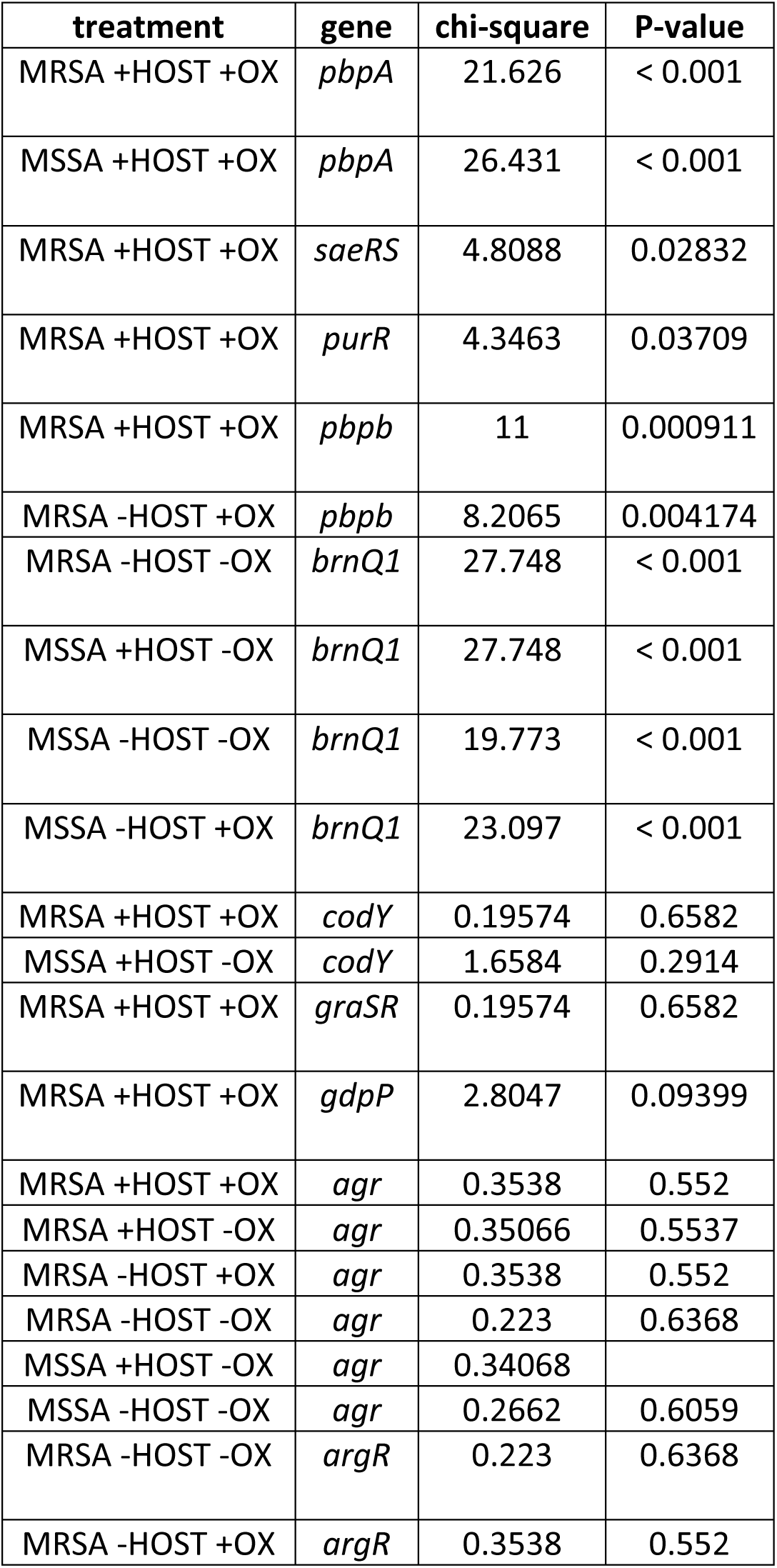
Chi-square goodness-of-fit test results for comparison between the proportion of blood/systemic infection-associated mutations *vs.* skin/nose/throat-associated mutations against the expected proportion (*i.e.,* all BioSamples in the dataset). All degrees of freedom equalled 1.

**Table S4.**
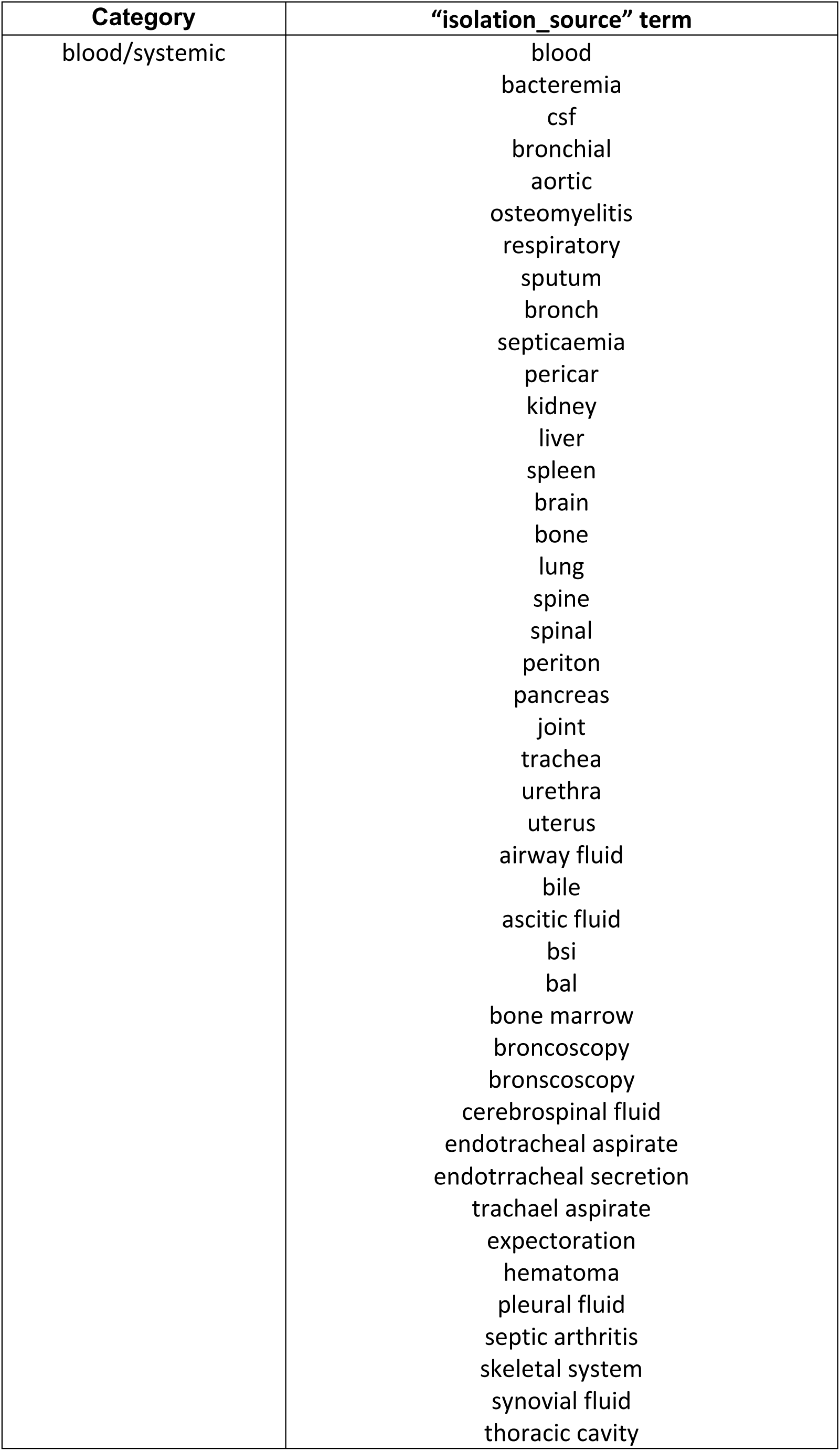
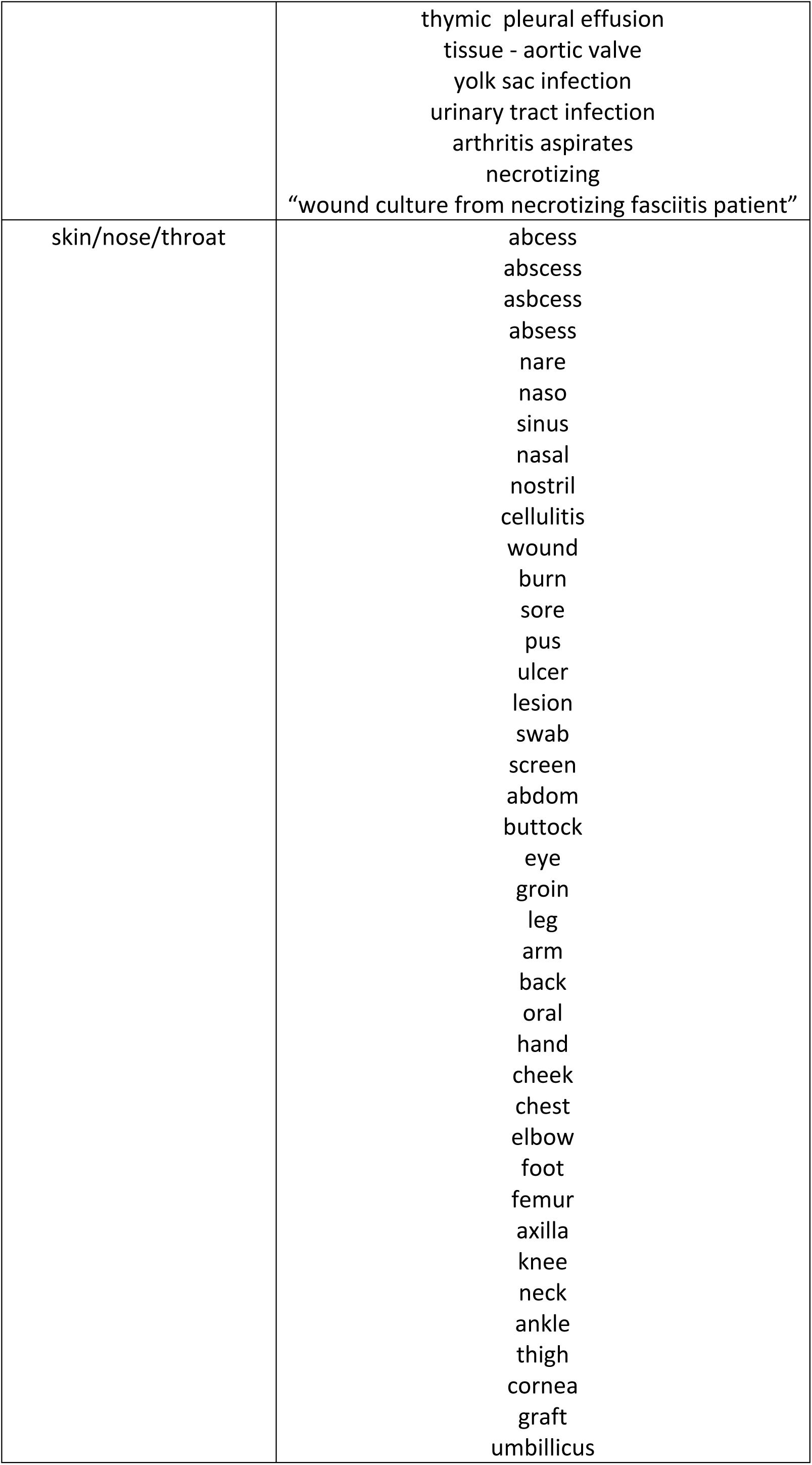
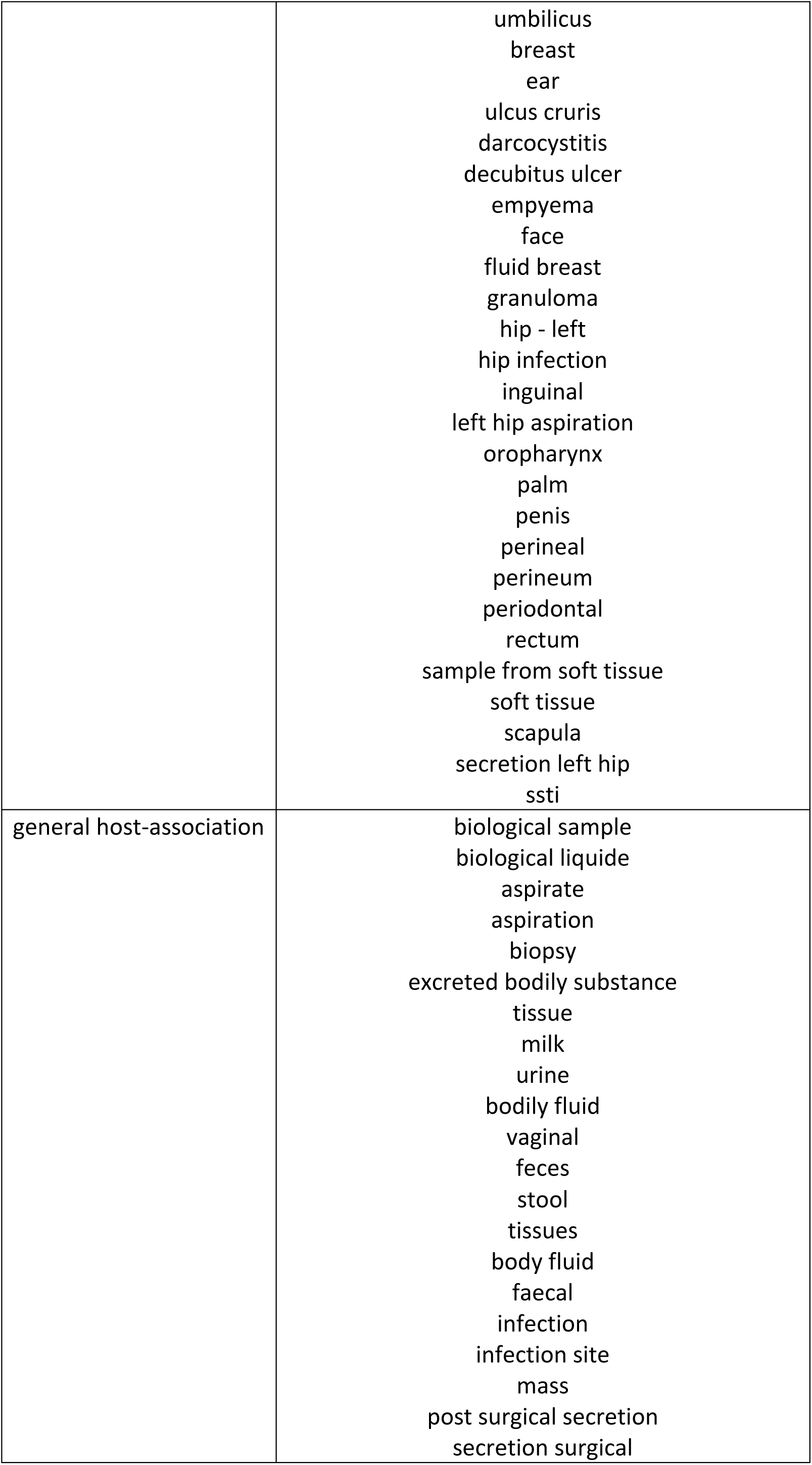
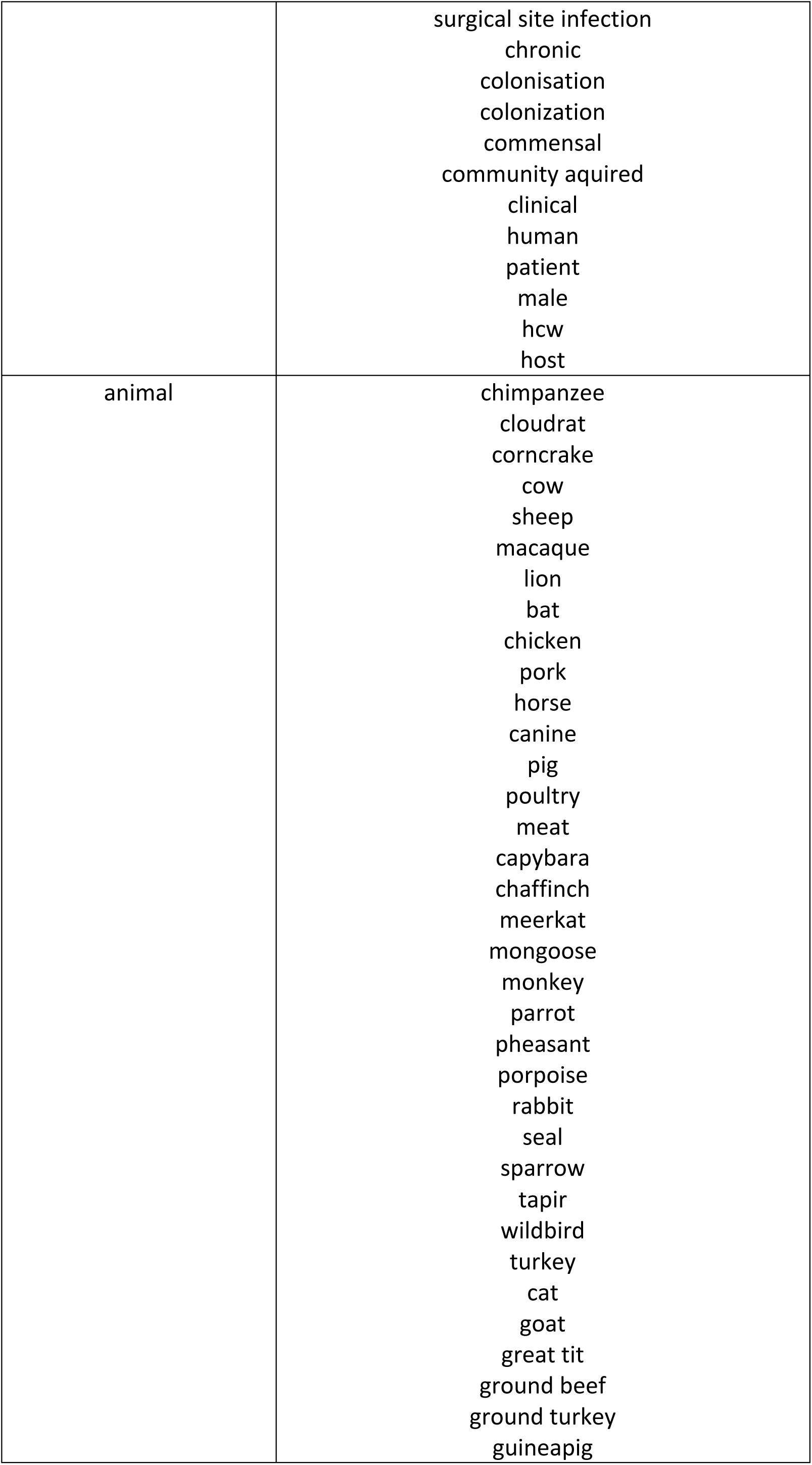
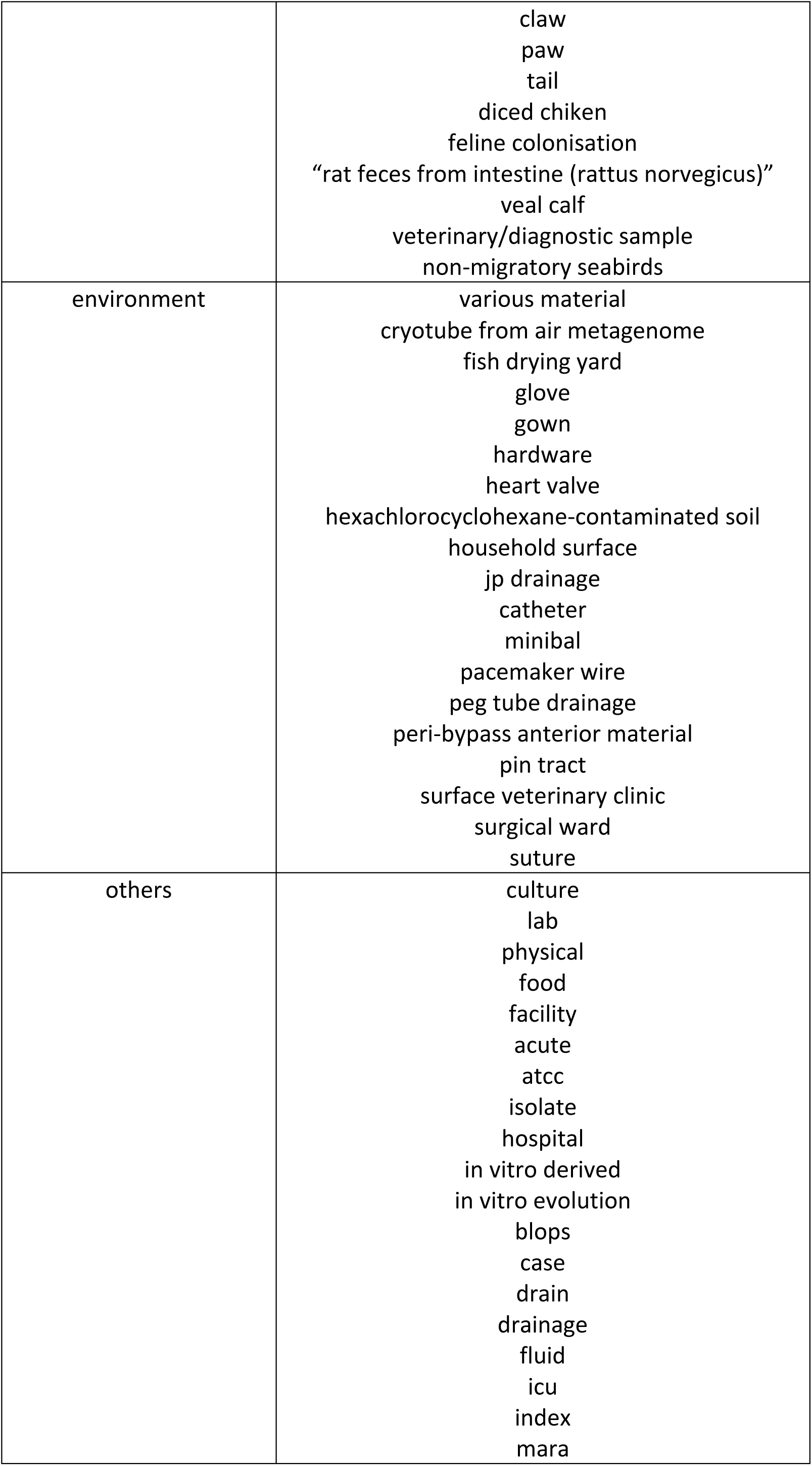
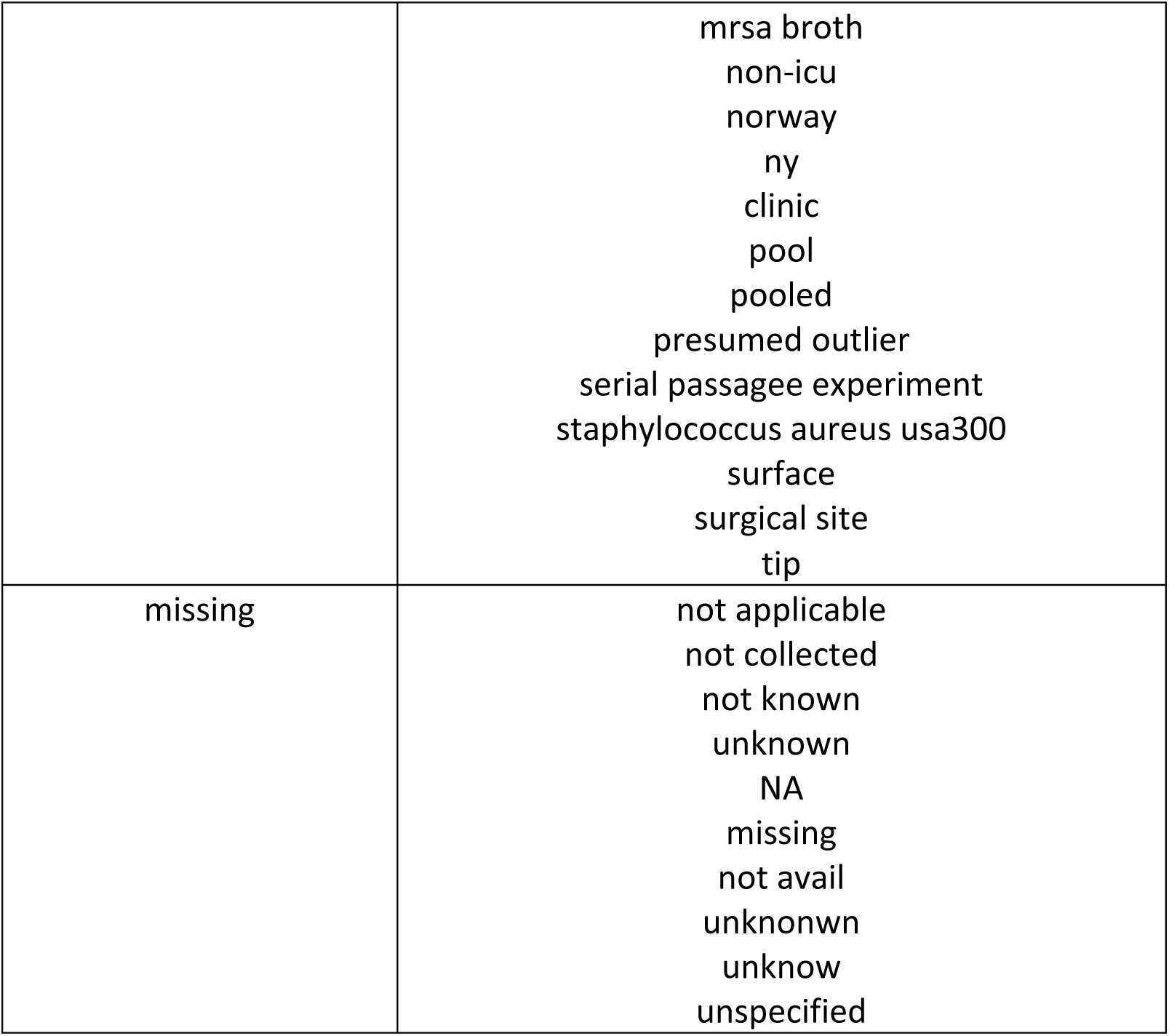
Categories for “isolation_source” terms from meta-data for BioSamples linked to our dataset of public *S. aureus* genomes in Figure 5A.

