## Supplemental Materials for "Host and antibiotic jointly select for greater virulence in *Staphylococcus aureus*"

**SUPPLEMENTAL FIGURES**

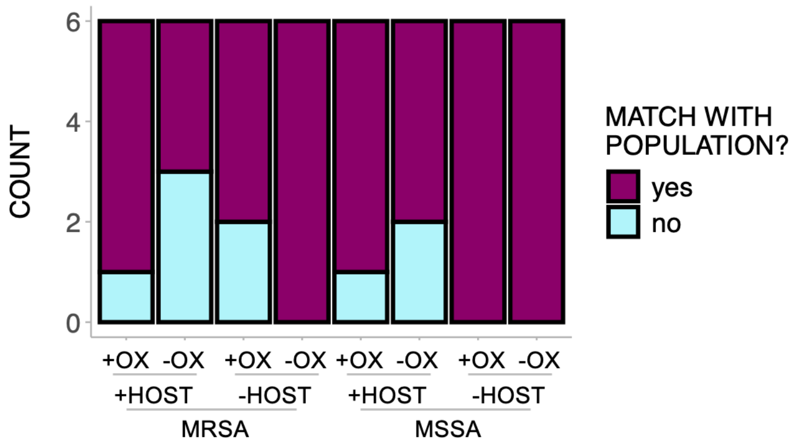


**Figure S1**. Most colonies sampled matched their respective population in terms of the ability to hemolyze sheep's blood (*i.e.,* over 50% of colonies having the same hemolysis status as the population they were sampled from).

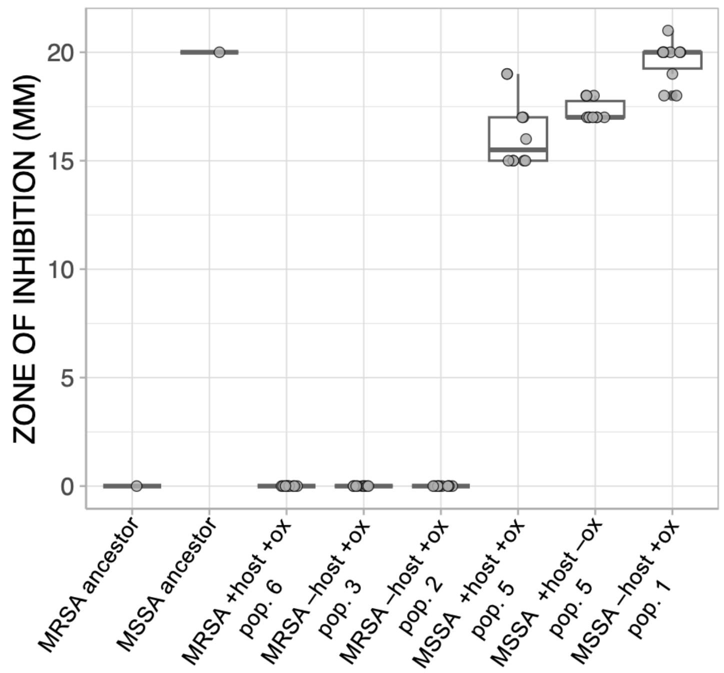


**Figure S2.** Zone of inhibition from Kirby-Bauer disk diffusion susceptibility test with oxacillin for colonies sampled from each of the four populations with the most number of mutations (two from MRSA and two from MSSA), and two additional randomly selected populations.

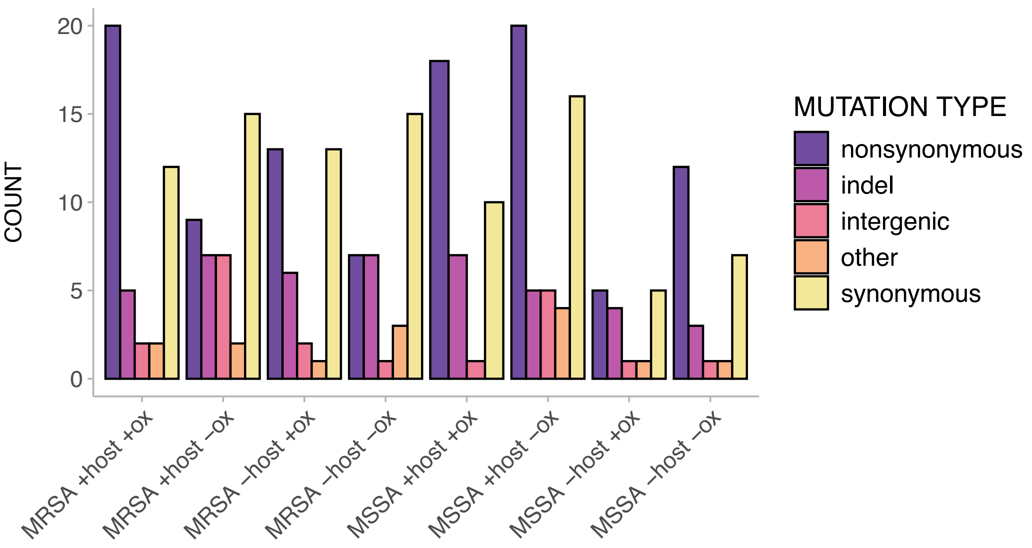


**Figure S3**. Count of all mutations (from 10-100% frequency) arisen in evolved populations.

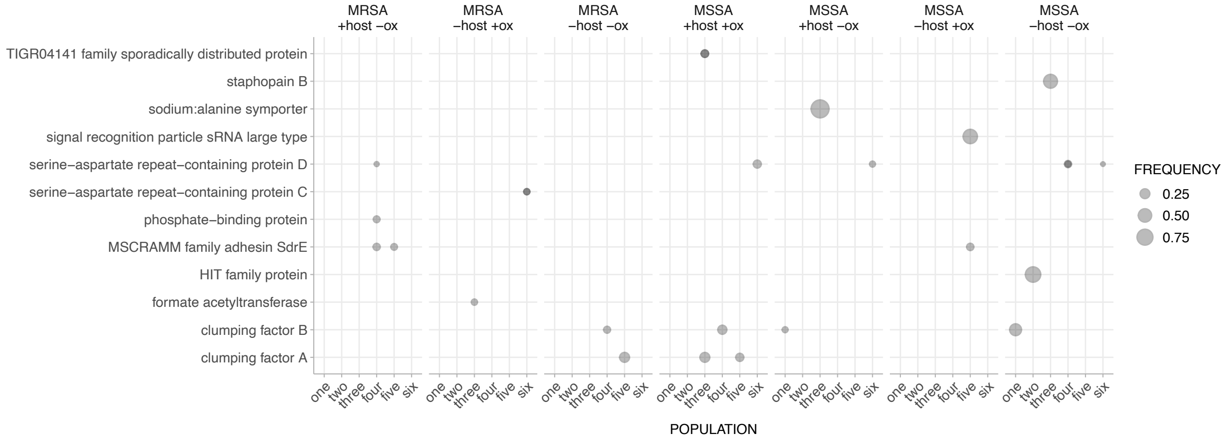


**Figure S4.** Mutations at frequencies between 0.1 and < 1, excluding synonymous or intergenic mutations, in each evolved population. Points with darker shades indicate more than one mutation present. All mutations in MRSA +host +ox populations between 0.1 and < 1 are all either synonymous or intergenic. A table of all mutations and frequencies is available on figshare (10.6084/m9.figshare.28745558)

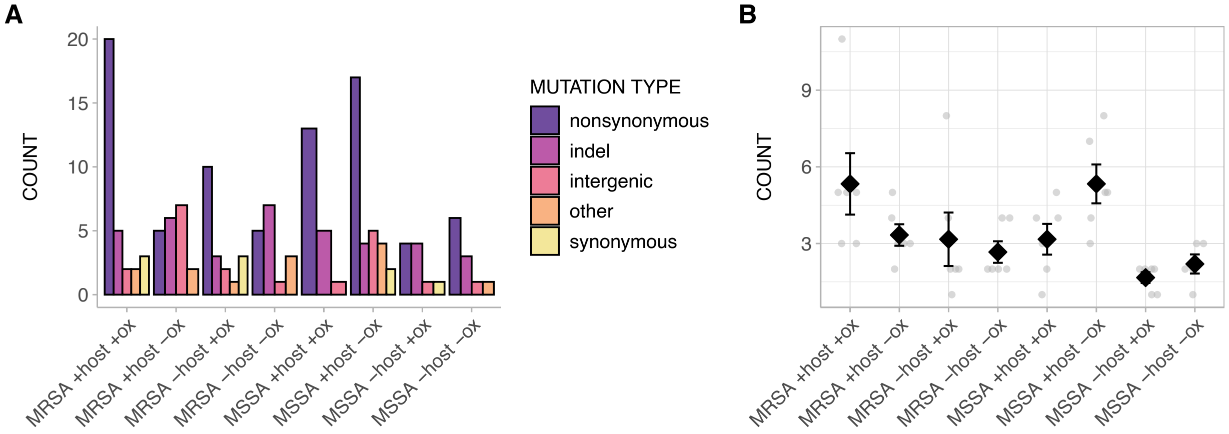


**Figure S5**. A) Count and B) Mean of mutations swept to fixation in evolved populations.

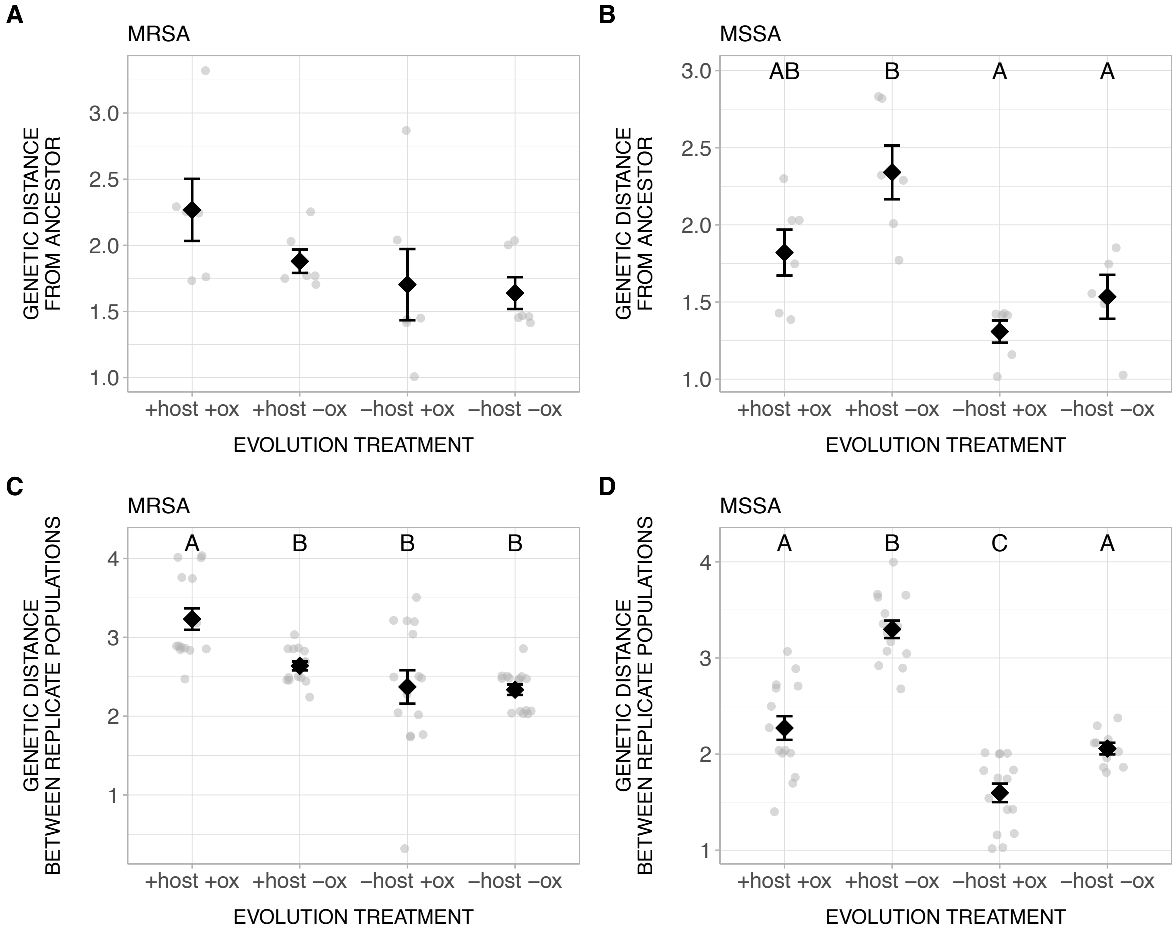


**Figure S6.** Genetic distance between the ancestor and evolved populations for A) MRSA and B) MSSA genotypes. Genetic distance between replicate populations within each treatment for C) MRSA and D) MSSA genotypes. Different letters indicate significant differences. Error bars indicate standard errors.

### SUPPLEMENTAL TABLES

**Table S1**. Nonsynonymous mutations occurred in genes with known roles in virulence and antibiotic resistance.

| Gene | Role identified in previous studies | Number of mutations in our study | Notes |
|---|---|---|---|
| *agr* | regulator, virulence [1] | 22 | 14 in MRSA<br>18 in -OX<br>10 in +HOST |
| *saeRS* | regulator, virulence [2] | 4 | 3 in MRSA |
| *codY* | regulator, metabolism and virulence [3] | 8 | 6 in MSSA<br>8 in +HOST<br>7 in +OX |
| *graSR* | regulator, virulence [4] and resistance to beta-lactam antibiotics [5] | 4 | 4 in MRSA<br>4 in +OX |
| *pbpA* | resistance to beta-lactam antibiotics [6] | 4 | 2 in MRSA<br>2 MSSA<br>4 in +OX |
| *pbpB* | resistance to beta-lactam antibiotics [7] | 4 | 4 MRSA<br>4 +OX |
| *gdpP* | resistance to beta-lactam antibiotics [8] | 4 | 1 in MRSA<br>3 in MSSA<br>4 in +OX |
| SCC*mec and* Arginine Catabolic Mobile Element (ACME) | resistance to beta-lactam antibiotics [9] | 9 | 5 SCC*mec* (23.7 kb deletion)<br>4 also included ACME (54.7 kb deletion)<br>6 in MSSA<br>7 in +HOST<br>1 *arcA* (99 bp deletion, encoded by ACME) in MRSA |
| *brnQ1* | metabolism [10] | 20 | 16 in -HOST |
| *purr* | virulence [11] | 4 | 4 in +HOST<br>3 in MRSA<br>3 in +OX |
| *gpmA* | virulence [12] | 2 | 2 in +HOST<br>2 in MSSA<br>2 in -OX |

**Table S2**. Fixed mutations in genes and intergenic regions appearing in more than two populations. SYN = synonymous, NONSYN = nonsynonymous

| Gene | Evolution treatment | Population | Position | Ancestral codon | Evolved codon | Mutation type | Mutation category | Amino acid change |
|---|---|---|---|---|---|---|---|---|
| ACME and *SCCmec* | MRSA -host -ox | three | 33633 | 54659-bp | | DEL | indel | |
| | MSSA +host +ox | three | 33633 | 54659-bp | | DEL | indel | |
| | MSSA +host -ox | five | 33633 | 54659-bp | | DEL | indel | |
| | | six | 33633 | 54659-bp | | DEL | indel | |
| B7H15_10520 | MRSA +host +ox | six | 2031543 | T | C | SNP | SYN | I19I |
| | MRSA -host +ox | three | 2031543 | T | C | SNP | SYN | I19I |
| *SCCmec* | MRSA +host -ox | three | 33636 | 23687-bp | | DEL | indel | |
| | | six | 33636 | 23687-bp | | DEL | indel | |
| | MSSA +host -ox | four | 33636 | 23687-bp | | DEL | indel | |
| | | five | 33636 | 23687-bp | | DEL | indel | |
| | MSSA -host +ox | two | 33636 | 23687-bp | | DEL | indel | |
| acetyl-CoA synthetase/ antibiotic biosynthesis monooxy-genase | MRSA +host -ox | two | 2749064 | C | A | SNP | intergenic | |
| | MSSA +host -ox | one | 2749081 | T | A | SNP | intergenic | |
| *agr* | MRSA +host +ox | six | 2151051 | C | T | SNP | NONSYN | S164L |
| | | six | 2151048 | A | G | SNP | NONSYN | E163G |
| | MRSA +host -ox | one | 2151213 | G | T | SNP | NONSYN | R218L |
| | | two | 2150580 | A | C | SNP | NONSYN | E7A |
| | | four | 2150063 | A | T | SNP | other | K256* |
| | MRSA +host -ox | five | 2151272 | A | | DEL | indel | |
| | | six | 2150931 | T | | DEL | indel | |
| | MRSA -host +ox | three | 2151051 | C | T | SNP | NONSYN | S164L |
| | | three | 2151048 | A | G | SNP | NONSYN | E163G |
| | MRSA -host -ox | two | 2149664 | A | | DEL | indel | |
| | | three | 2151047 | G | T | SNP | other | E163* |
| | | four | 2151272 | A | | DEL | indel | |
| | | five | 2149476 | G | A | SNP | other | W60* |
| | | six | 2151212 | C | T | SNP | other | R218* |
| | MSSA +host -ox | four | 2150250 | G | A | SNP | NONSYN | G318D |
| | | five | 2151233 | A | T | SNP | other | K225* |
| | | six | 2150298 | C | A | SNP | NONSYN | P334H |
| | MSSA -host -ox | one | 2149635 | C | A | SNP | other | S113* |
| | | three | 2150803 | T | CAGTT | INS | indel | |
| | | four | 2149989 | C | A | SNP | NONSYN | T231K |
| | | six | 2150700 | C | A | SNP | NONSYN | A47D |
| | | six | 2150691 | TT | AA | SUB | indel | |

| | | | | | | | | |
|---|---|---|---|---|---|---|---|---|
| *agrD/C* | MRSA -host +ox | one | 2149252 | G | A | SNP | intergenic | |
| | MSSA -host -ox | two | 2149281 | T | G | SNP | intergenic | |
| alr | MRSA -host +ox | two | 2190276 | A | G | SNP | NONSYN | L280S |
| | | six | 2190276 | A | G | SNP | NONSYN | L280S |
| | MRSA -host -ox | one | 2190276 | A | G | SNP | NONSYN | L280S |
| *argR* | MRSA +host +ox | four | 1639665 | G | C | SNP | NONSYN | T99R |
| | MRSA -host +ox | two | 1639510 | A | T | SNP | NONSYN | *151K |
| | | six | 1639510 | A | T | SNP | NONSYN | *151K |
| | MRSA -host -ox | one | 1639510 | A | T | SNP | NONSYN | *151K |
| arginine repressor/-geranyl transferase | MRSA +host -ox | three | 1639991 | C | T | SNP | intergenic | |
| | MSSA +host -ox | four | 1639991 | C | T | SNP | intergenic | |
| | | six | 1639991 | C | T | SNP | intergenic | |
| *brnQ1* | MRSA -host -ox | two | 218284 | A | G | SNP | NONSYN | L154S |
| | | three | 218456 | A | T | SNP | NONSYN | F97I |
| | | three | 218451 | G | | DEL | indel | |
| | | four | 218425 | 108-bp | | DEL | indel | |
| | | five | 217434 | GCCGATAAT GAA | | DEL | indel | |
| | | six | 217538 | 97-bp | | DEL | indel | |
| | MSSA +host +ox | three | 218420 | CA | AT | SUB | indel | |
| | MSSA +host -ox | two | 218581 | A | T | SNP | other | L55* |
| | | three | 217879 | G | T | SNP | other | S289* |
| | | five | 218644 | C | A | SNP | NONSYN | G34V |
| | MSSA -host +ox | one | 218035 | C | T | SNP | NONSYN | G237D |
| | | two | 218420 | CA | AT | SUB | indel | |
| | | three | 218420 | CA | AT | SUB | indel | |
| | | four | 218035 | C | T | SNP | NONSYN | G237D |
| | | five | 218420 | CA | AT | SUB | indel | |
| | | six | 218035 | C | T | SNP | NONSYN | G237D |
| | MSSA -host -ox | one | 218128 | GCTAGCGCA T | | DEL | indel | |
| | | two | 218644 | C | A | SNP | NONSYN | G34V |
| | | three | 218644 | C | A | SNP | NONSYN | G34V |
| | | six | 218644 | C | A | SNP | NONSYN | G34V |
| *codY* | MRSA +host +ox | two | 1275011 | C | T | SNP | NONSYN | H159Y |
| | | six | 1274675 | A | G | SNP | NONSYN | K47E |
| | | one | 1275155 | G | T | SNP | NONSYN | A207S |
| | | two | 1275155 | G | T | SNP | NONSYN | A207S |
| | MSSA +host +ox | four | 1275155 | G | T | SNP | NONSYN | A207S |
| | | five | 1275155 | G | T | SNP | NONSYN | A207S |
| | | six | 1275155 | G | T | SNP | NONSYN | A207S |
| | MSSA +host -ox | two | 1274976 | C | T | SNP | NONSYN | A147V |
| | MRSA +host +ox | six | 87329 | T | A | SNP | SYN | L624L |

| | | | | | | | | |
|---|---|---|---|---|---|---|---|---|
| copper-translo-cating P-type ATPase | MRSA -host +ox | three | 87329 | T | A | SNP | SYN | L624L |
| *ebh* | MSSA +host -ox | five | 1477894 | C | T | SNP | NONSYN | E8357K |
| | MSSA -host -ox | two | 1480506 | G | A | SNP | NONSYN | P7486L |
| *fmtB* | MRSA +host +ox | six | 2284788 | G | T | SNP | other | |
| | MRSA -host +ox | three | 2284788 | G | T | SNP | other | |
| *gdpP* | MRSA +host +ox | three | 18721 | G | T | SNP | NONSYN | G307V |
| | MSSA +host +ox | one | 18720 | G | A | SNP | NONSYN | G307S |
| | | four | 18745 | A | C | SNP | NONSYN | K315T |
| | | five | 19143 | A | T | SNP | NONSYN | S448C |
| *gpmA* | MSSA +host -ox | one | 2539086 | G | A | SNP | NONSYN | R114C |
| | MSSA +host -ox | three | 2538887 | G | A | SNP | NONSYN | S180F |
| *graSR* | MRSA +host +ox | three | 734510 | A | | DEL | indel | |
| | | four | 735131 | C | G | SNP | other | S187* |
| | | six | 734085 | T | | DEL | indel | |
| | MRSA -host +ox | three | 734085 | T | | DEL | indel | |
| Hydroxy-methyl-glutaryl-CoA synthase/-cysteine methyl-transferase | MRSA +host +ox | six | 2685330 | T | C | SNP | intergenic | |
| | MRSA -host +ox | three | 2685330 | T | C | SNP | intergenic | |
| *pbpA* | MRSA +host +ox | one | 1189756 | C | A | SNP | NONSYN | D350E |
| | | four | 1190201 | C | G | SNP | NONSYN | H499D |
| | MSSA +host +ox | three | 1189756 | C | A | SNP | NONSYN | D350E |
| | | six | 1189758 | G | T | SNP | NONSYN | W351L |
| *pbpB* | MRSA +host +ox | two | 1520209 | G | A | SNP | NONSYN | A132T |
| | | five | 1520434 | G | A | SNP | NONSYN | A207T |
| | MRSA -host +ox | one | 1520247 | A | GGTTCTG AA | INS | indel | |
| | | four | 1520414 | G | T | SNP | NONSYN | G200V |
| *purR* | MRSA +host +ox | one | 532212 | C | T | SNP | NONSYN | R96C |
| | | three | 532013 | T | T | INS | indel | |
| | | five | 532013 | T | T | INS | indel | |
| | MRSA -host +ox | four | 532240 | T | T | INS | indel | |
| *saeRS* | MRSA +host +ox | two | 782595 | G | T | SNP | NONSYN | P74T |
| | | five | 782577 | C | T | SNP | NONSYN | A80T |
| | MRSA +host -ox | two | 781408 | G | A | SNP | NONSYN | R241C |
| | MSSA +host +ox | six | 782211 | C | | DEL | indel | |

**Table S3.** Chi-square goodness-of-fit test results for comparison between the proportion of blood/systemic infection-associated mutations *vs.* skin/nose/throat-associated mutations against the expected proportion (*i.e.,* all BioSamples in the dataset). All degrees of freedom equalled 1.

| treatment | gene | chi-square | P-value |
|---|---|---|---|
| MRSA +HOST +OX | *pbpA* | 21.626 | < 0.001 |
| MSSA +HOST +OX | *pbpA* | 26.431 | < 0.001 |
| MRSA +HOST +OX | *saeRS* | 4.8088 | 0.02832 |
| MRSA +HOST +OX | *purR* | 4.3463 | 0.03709 |
| MRSA +HOST +OX | *pbpb* | 11 | 0.000911 |
| MRSA -HOST +OX | *pbpb* | 8.2065 | 0.004174 |
| MRSA -HOST -OX | *brnQ1* | 27.748 | < 0.001 |
| MSSA +HOST -OX | *brnQ1* | 27.748 | < 0.001 |
| MSSA -HOST -OX | *brnQ1* | 19.773 | < 0.001 |
| MSSA -HOST +OX | *brnQ1* | 23.097 | < 0.001 |
| MRSA +HOST +OX | *codY* | 0.19574 | 0.6582 |
| MSSA +HOST -OX | *codY* | 1.6584 | 0.2914 |
| MRSA +HOST +OX | *graSR* | 0.19574 | 0.6582 |
| MRSA +HOST +OX | *gdpP* | 2.8047 | 0.09399 |
| MRSA +HOST +OX | *agr* | 0.3538 | 0.552 |
| MRSA +HOST -OX | *agr* | 0.35066 | 0.5537 |
| MRSA -HOST +OX | *agr* | 0.3538 | 0.552 |
| MRSA -HOST -OX | *agr* | 0.223 | 0.6368 |
| MSSA +HOST -OX | *agr* | 0.34068 | |
| MSSA -HOST -OX | *agr* | 0.2662 | 0.6059 |
| MRSA -HOST -OX | *argR* | 0.223 | 0.6368 |
| MRSA -HOST +OX | *argR* | 0.3538 | 0.552 |

**Table S4.** Categories for “isolation_source” terms from meta-data for BioSamples linked to our dataset of public *S. aureus* genomes in Figure 5A.

| Category | “isolation_source” term |
|---|---|
| blood/systemic | blood |
| | bacteremia |
| | csf |
| | bronchial |
| | aortic |
| | osteomyelitis |
| | respiratory |
| | sputum |
| | bronch |
| | septicaemia |
| | pericar |
| | kidney |
| | liver |
| | spleen |
| | brain |
| | bone |
| | lung |
| | spine |
| | spinal |
| | periton |
| | pancreas |
| | joint |
| | trachea |
| | urethra |
| | uterus |
| | airway fluid |
| | bile |
| | ascitic fluid |
| | bsi |
| | bal |
| | bone marrow |
| | broncoscopy |
| | bronscoscopy |
| | cerebrospinal fluid |
| | endotracheal aspirate |
| | endotrracheal secretion |
| | trachael aspirate |
| | expectoration |
| | hematoma |
| | pleural fluid |
| | septic arthritis |
| | skeletal system |
| | synovial fluid |
| | thoracic cavity |

| | |
|---|---|
| | thymic pleural effusion<br>tissue - aortic valve<br>yolk sac infection<br>urinary tract infection<br>arthritis aspirates<br>necrotizing<br>“wound culture from necrotizing fasciitis patient” |
| skin/nose/throat | abcess<br>abscess<br>asbcess<br>absess<br>nare<br>naso<br>sinus<br>nasal<br>nostril<br>cellulitis<br>wound<br>burn<br>sore<br>pus<br>ulcer<br>lesion<br>swab<br>screen<br>abdom<br>buttock<br>eye<br>groin<br>leg<br>arm<br>back<br>oral<br>hand<br>cheek<br>chest<br>elbow<br>foot<br>femur<br>axilla<br>knee<br>neck<br>ankle<br>thigh<br>cornea<br>graft<br>umbillicus |

| | |
|---|---|
| | umbilicus<br>breast<br>ear<br>ulcus cruris<br>darcocystitis<br>decubitus ulcer<br>empyema<br>face<br>fluid breast<br>granuloma<br>hip - left<br>hip infection<br>inguinal<br>left hip aspiration<br>oropharynx<br>palm<br>penis<br>perineal<br>perineum<br>periodontal<br>rectum<br>sample from soft tissue<br>soft tissue<br>scapula<br>secretion left hip<br>ssti |
| general host-association | biological sample<br>biological liquide<br>aspirate<br>aspiration<br>biopsy<br>excreted bodily substance<br>tissue<br>milk<br>urine<br>bodily fluid<br>vaginal<br>feces<br>stool<br>tissues<br>body fluid<br>faecal<br>infection<br>infection site<br>mass<br>post surgical secretion<br>secretion surgical |

| | |
|---|---|
| | surgical site infection<br>chronic<br>colonisation<br>colonization<br>commensal<br>community aquired<br>clinical<br>human<br>patient<br>male<br>hcw<br>host |
| animal | chimpanzee<br>cloudrat<br>corncrake<br>cow<br>sheep<br>macaque<br>lion<br>bat<br>chicken<br>pork<br>horse<br>canine<br>pig<br>poultry<br>meat<br>capybara<br>chaffinch<br>meerkat<br>mongoose<br>monkey<br>parrot<br>pheasant<br>porpoise<br>rabbit<br>seal<br>sparrow<br>tapir<br>wildbird<br>turkey<br>cat<br>goat<br>great tit<br>ground beef<br>ground turkey<br>guineapig |

| | |
|---|---|
| | claw<br>paw<br>tail<br>diced chiken<br>feline colonisation<br>“rat feces from intestine (rattus norvegicus)”<br>veal calf<br>veterinary/diagnostic sample<br>non-migratory seabirds |
| environment | various material<br>cryotube from air metagenome<br>fish drying yard<br>glove<br>gown<br>hardware<br>heart valve<br>hexachlorocyclohexane-contaminated soil<br>household surface<br>jp drainage<br>catheter<br>minibal<br>pacemaker wire<br>peg tube drainage<br>peri-bypass anterior material<br>pin tract<br>surface veterinary clinic<br>surgical ward<br>suture |
| others | culture<br>lab<br>physical<br>food<br>facility<br>acute<br>atcc<br>isolate<br>hospital<br>in vitro derived<br>in vitro evolution<br>blops<br>case<br>drain<br>drainage<br>fluid<br>icu<br>index<br>mara |

| | |
|---|---|
| | mrsa broth<br>non-icu<br>norway<br>ny<br>clinic<br>pool<br>pooled<br>presumed outlier<br>serial passagee experiment<br>staphylococcus aureus usa300<br>surface<br>surgical site<br>tip |
| missing | not applicable<br>not collected<br>not known<br>unknown<br>NA<br>missing<br>not avail<br>unknonwn<br>unknow<br>unspecified |
